# Prefrontal cortex glucocorticoid receptors during fear memory consolidation shift the balance between salience and default-mode networks at retrieval in rats

**DOI:** 10.64898/2026.08.21.745892

**Authors:** Moisés dos Santos Corrêa, Lorena Vido Lopes, Ana Carolina Quintiliano dos Santos, Juliana Camino Castro, William Thiago Boscariol Lourenço, Ana Carolina da Costa Silva, Tatiana Lima Ferreira, Paula Ayako Tiba, Raquel Vecchio Fornari

## Abstract

Contextual fear memories can lose specificity as they age, modeling fear overgeneralization seen in post-traumatic stress disorder. Glucocorticoid receptor (GR) signaling in the dorsomedial prefrontal cortex (dmPFC) during the immediate post-learning period may govern both endocrine recovery from an aversive experience and the eventual specificity of the resulting memory, but this link remains untested. We infused vehicle or the GR antagonist mifepristone into the dmPFC of rats immediately after contextual fear conditioning, then assessed corticosterone dynamics, behavior and c-Fos coactivation networks at recent and remote time points. Mifepristone accelerated corticosterone recovery without changing hormone release, spared recent memory, and produced stronger, generalized defensive behavior at the remote time point. This behavioral shift coincided with denser remote retrieval networks and greater default-mode-like than salience-like connectivity. These findings identify post-learning dmPFC GR signaling as a regulator of endocrine recovery and remote defensive expression, constraining fear memory specificity as memories undergo systems consolidation.

## INTRODUCTION

Contextual fear memories change qualitatively as they age. Hippocampal-dependent representations initially allow precise discrimination between dangerous and safe contexts (Anagnostaras et al., 2001; Asok et al., 2018; Atucha et al., 2017; de Oliveira Alvares & Do-Monte, 2021; Frankland et al., 1998), but these representations undergo systems consolidation over subsequent days to weeks, progressively engaging neocortical regions including the medial prefrontal cortex (mPFC) and, in some cases, losing contextual specificity in the process (Dos Santos Corrêa et al., 2019; Frankland & Bontempi, 2005; Lux et al., 2016; Maviel et al., 2004; Pedraza et al., 2016; Quillfeldt, 2019). This strengthening of fear memory and loss of specificity, or time-dependent fear generalization, resembles a feature of post-traumatic stress disorder (PTSD) and related anxiety disorders, in which patients experience heightened and inappropriate fear responses to safe cues or contexts that merely resemble a traumatic one (Kaczkurkin et al., 2017; Lissek et al., 2005; Webler et al., 2021). Glucocorticoids, released by the hypothalamic-pituitary-adrenal (HPA) axis, exert time-dependent control over this process (Barsegyan et al., 2010; Roozendaal, 2002). Glucocorticoid receptor (GR) activation around the time of learning enhances memory consolidation (Roozendaal, 1999), and within the mPFC specifically, GR signaling both facilitates consolidation (Roozendaal et al., 2009) and provides negative feedback that terminates the HPA stress response (Hill et al., 2011; McKlveen et al., 2013; Radley, 2012), making this region a convergence point between the endocrine and mnemonic consequences of an aversive experience. Within the mPFC, the prelimbic (PrL) and anterior cingulate (ACC) subdivisions, together referred to here as the dorsomedial prefrontal cortex (dmPFC), are implicated in modulating the strength and precision of contextual fear memory at recent time points as well as it transitions to a remote form (Bian et al., 2019; Frankland et al., 2004; Restivo et al., 2009; Sierra et al., 2017; Stern et al., 2014; Vanvossen et al., 2017), although PrL and ACC are not functionally interchangeable and can contribute differentially to fear expression (Bayer et al., 2022; da Silva et al., 2020; Santos et al., 2017; Troyner et al., 2018). Whether GR signaling in the dmPFC during the immediate post-learning window shapes both the endocrine trajectory of the stress response and the eventual strength or specificity of the resulting memory remains unknown.

A further gap concerns how glucocorticoid-dependent processes in the dmPFC shape the broader brain network supporting contextual fear memory. Because fear memories are not stored in a single region but are distributed across interacting hippocampal, amygdalar, and cortical regions (Dixsaut & Gräff, 2021; Roy et al., 2022; Vetere et al., 2017), understanding their reorganization over time requires network-level measures, such as graph-theoretic metrics of regional hub status and community structure, rather than activity changes in individual regions alone; such reorganization can be traced using c-Fos coactivation across this network (Dos Santos Corrêa et al., 2022; Santos et al., 2023). In humans, acute stress initially favors a salience network (SN) configuration, including the amygdala and dmPFC, while the delayed, corticosteroid-dependent phase of the stress response favors a default-mode network (DMN)-like configuration instead (Schwabe et al., 2022), consistent with a broader role for GR in shifting the brain from a vigilance-oriented to an integrative, consolidation-oriented mode (de Kloet & Joëls, 2023). A recent study showed that mice susceptible to stress vulnerability have increased SN-DMN coupling around and after severe stress exposure compared to resilient mice (Dirven et al., 2026). We reasoned that GR signaling in the dmPFC during post-training consolidation may similarly influence the balance between SN-like and DMN-like states within the fear memory network, with consequences for fear memory strength and expression at remote time points.

Prior work provides indirect support for a link between dmPFC glucocorticoid signaling and the fate of contextual fear memories. Post-training manipulations of rostral PrL projections to the anteroventral bed nucleus of the stria terminalis also bidirectionally regulate contextual discrimination even in the absence of stress-evoked CORT release (Lingg et al., 2026). Glucocorticoid administration into the mPFC after learning enhances memory consolidation (Barsegyan et al., 2010) while impairing working memory (Butts et al., 2011), indicating that this region is sensitive to glucocorticoid levels during a defined post-learning window. Separately, glucocorticoid signaling in the dentate gyrus and lateral amygdala biases engram allocation toward broader, less selective ensembles, a mechanism proposed to underlie stress-induced fear generalization (Lesuis et al., 2021, 2025). The dmPFC already allocates a subset of neurons to the fear engram during acquisition itself, with this ensemble later becoming functionally engaged only as the memory transitions to a remote, cortically-dependent state (Matos et al., 2019). We reasoned that GR signaling in the dmPFC during the immediate post-training period may act on this allocated ensemble by shaping its later contribution to memory expression. Blocking dmPFC GRs after training may therefore affect both the endocrine recovery from the aversive experience and the strength of the memory at remote retrieval. More broadly, an imbalance in MR-and GR-mediated actions has been proposed to compromise adaptive stress management and promote vulnerability to stress-related psychopathology (de Kloet & Joëls, 2023), raising the possibility that dmPFC GR signaling during consolidation similarly determines whether an animal remains resilient to, or becomes vulnerable to, altered fear memory strength and expression as the memory ages.

To address this, we implanted bilateral cannulae targeting the dmPFC in male rats and infused vehicle (VEH) or the GR antagonist mifepristone (MIF) immediately after contextual fear conditioning (CFC). We hypothesized that dmPFC GR signaling during consolidation regulates the balance between SN-like and DMN-like network configurations during early consolidation, thereby determining an animal’s resilience or vulnerability to altered fear memory strength and expression at remote time points. To test this, we tracked post-training CORT dynamics, assessed fear memory strength and specificity at recent (2-day) and remote (14-day) time points, and used c-Fos coactivation networks to determine whether MIF-induced shifts in endocrine recovery and fear memory strength were paralleled by reorganization of SN-like and DMN-like retrieval networks.

## RESULTS

### Post-training dmPFC GR blockade alters CORT dynamics during contextual fear memory consolidation

The mPFC, particularly its PrL and IL subdivisions, has been implicated in glucocorticoid-mediated negative feedback control of the HPA axis during emotional stress (Herman et al., 2016; Radley, 2012; Ulrich-Lai & Herman, 2009). Based on this, we tested whether blocking GRs in the dmPFC immediately after CFC would interfere with post-training CORT feedback regulation, thereby altering the endocrine recovery that normally follows aversive learning (Cordero & Sandi, 1998; McKlveen et al., 2013).

Cannula-implanted male rats underwent CFC followed immediately by bilateral infusions of VEH or MIF into the dmPFC; tail blood samples were then collected 30 and 60 min after infusions for CORT quantification (**Figure 1A**). 46 implanted rats were used, of which 3 were excluded after histological analysis supporting anatomical specificity of the infusion procedure (cannula tips mainly in the PrL and some in the ACC), leaving 43 for analysis (**Figure 1B**). Rats from both treatment groups (VEH n = 20, MIF n = 23) increased freezing during the post-shock interval of training compared to the pre-shocks interval, indicating there was no intrinsic bias in either group towards innate high baseline freezing (**Figure 1C**).

**Figure 1.**
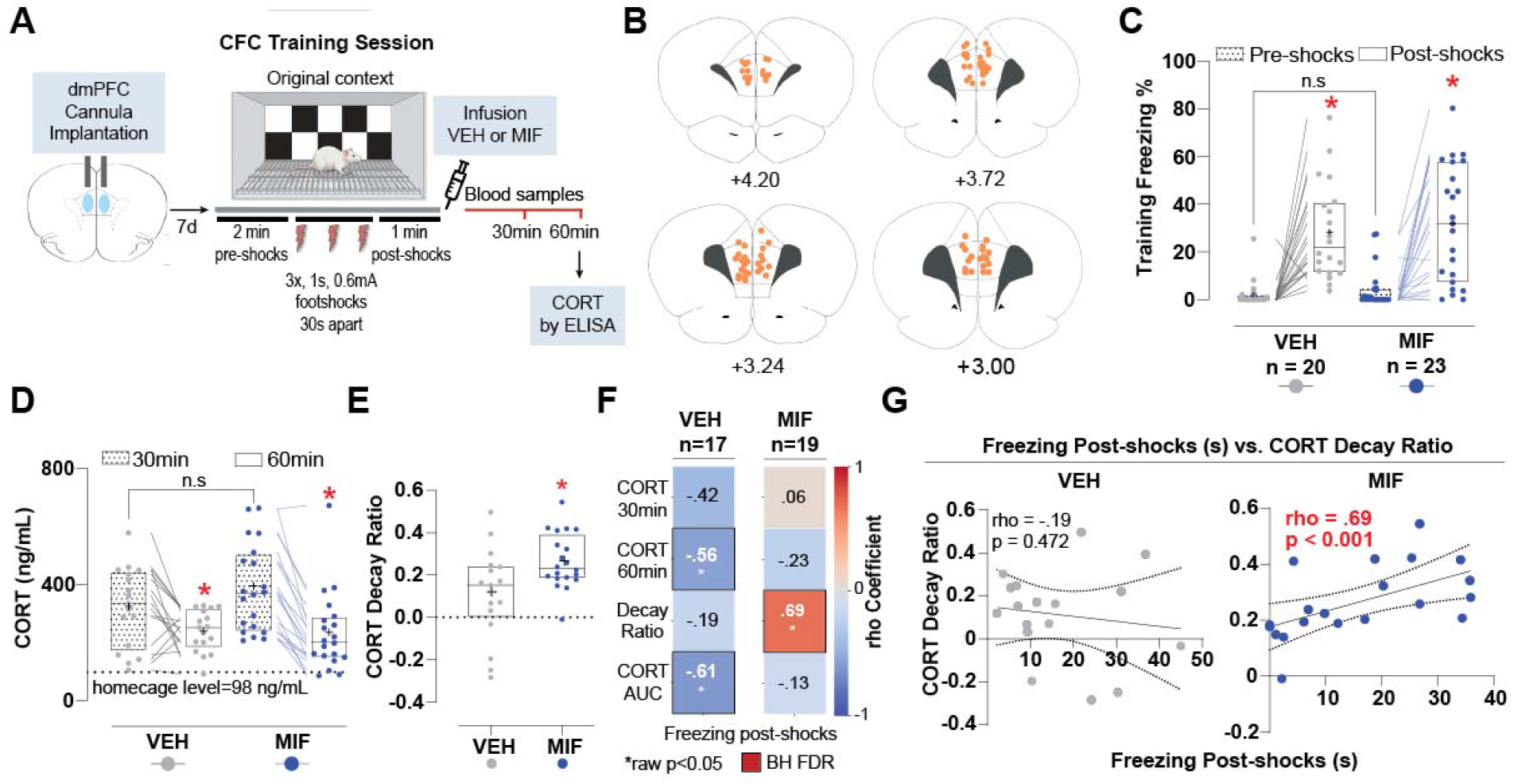
Post-training dmPFC GR blockade alters CORT dynamics during contextual fear memory consolidation. **(A)** Schematic of experimental design. Rats were implanted with bilateral guide cannulae targeting the dorsomedial prefrontal cortex (dmPFC), allowed to recover, and then subjected to a contextual fear conditioning (CFC) task consisting of three footshocks (0.6 mA, 1 s) in an unfamiliar training context. Immediately after training, rats received intra-dmPFC infusions of vehicle (VEH) or the GR antagonist mifepristone (MIF), followed by tail blood collection at 30 and 60 min after infusion ending for corticosterone (CORT) quantification by ELISA. **(B)** Verification of dmPFC cannula placements. Coronal schematics depict the distribution of injector tips (orange circles) across anterior–posterior levels relative to bregma in rats included in the analyses. **(C-D)** Behavioral and endocrine repeated measurements referring to the training session. Datapoints from both groups (VEH and MIF) are shown in different colors (gray and blue respectively). Lines connect the different repeated measures values for individual animals, and boxplots showing median and interquartile intervals. The plus sign reflects the mean for each subset of data. For both panels, statistical differences were quantified using Mixed ANOVAs, p-values < 0.05 were considered significant, n.s. for non-significant comparison. **(C)** Median freezing percentage during the training session ± IQR. Freezing was measured during the pre-shock and post-shocks periods of CFC training. Mixed ANOVA: Freezing F(1,41) = 37.41, p < 0.001, η²_G_ = 0.28; Treatment F(1,41) = 0.83, p = 0.37, η²_G_ = 0.01; Freezing*Treatment F(1,41) = 0.01, p = 0.92, η²_G_ = 0.00. **(D)** Median plasma CORT concentrations ± IQR measured 30 and 60 min after post-training treatment are shown for VEH- and MIF-treated rats, with the homecage baseline level indicated by a dashed line. Mixed ANOVA: Concentration F(1,35) = 47.32, p < 0.001, η²_G_ = 0.16; Treatment F(1,35) = 0.55, p = 0.46, η²_G_ = 0.01; Concentration*Treatment F(1,35) = 4.14, p = 0.05, η²_G_ = 0.02. **(E)** Median ratio of CORT decay ± IQR between 30 and 60 min after training for VEH (gray) or MIF (blue). CORT decay ratio reflects the proportional decline in corticosterone from the 30-to 60-minute post-stress sample, calculated as (CORT30min − CORT60min)/(CORT30min + CORT60min). Higher values reflect faster hormonal recovery, while values around zero stability over the time interval. 2 samples Welch’s *t* test: t(35) = 2.68, p = 0.01, d = 0.88. *p-value < 0.05 when comparing to the VEH group. **(F)** Heatmaps with Spearman correlation coefficients (rho) between Freezing post-shocks and CORT measures (30 min, 60 min, decay ratio, and AUC) in VEH-(left) and MIF-treated (right) rats. Color scale indicates direction and magnitude of the correlations, with asterisks denoting coefficients that reach statistical significance and bold square borders indicating correlations that remain significant after BH FDR correction for multiple comparisons. **(G)** Targeted correlations linking post-shocks freezing response to CORT decay ratio in VEH (left) and MIF (right) groups, after Holm correction. Each scatterplot shows data from individual animals with linear regression fits and 95% confidence intervals. Statistical results (rho and p values) are shown within each panel.

Post-training CORT plasma levels for both treatment groups (VEH n = 17, MIF n = 20) remained elevated above the indicated home-cage group baseline (average 98 ng/mL, n = 6) at both 30 and 60 min after contextual fear conditioning, with CORT levels at 60 min being significantly lower than those at 30 min (**Figure 1D**). Additionally, total released CORT, calculated as CORT area under the curve (AUC) across the 30-60 min interval, was not significantly different between groups (**Supplementary Figure 1A**). On the other hand, MIF-treated rats showed higher Decay Ratio compared to the VEH group, suggesting that the treatment influenced the endocrine recovery (**Figure 1E**).

Correlation analyses further showed that the relationship between time spent in freezing during the post-shock interval of the training and CORT response differed by treatment condition, with a distinct association between freezing and CORT Decay Ratio across groups (**Figure 1F**). Specifically, in the MIF group, post-shock freezing levels correlated positively with the CORT Decay Ratio, indicating that under MIF treatment, stronger fear expression during training predicted a faster endocrine recovery during consolidation (**Figure 1G**). Together, these findings indicate that dmPFC GR blockade reshapes the dynamics linking fear conditioning to glucocorticoid response during memory consolidation. Given this, we next asked whether this altered endocrine trajectory during consolidation translates into changes in fear memory strength and generalization during retrieval.

### dmPFC GR blockade during consolidation spares recent but alters remote fear memory

Prior work shows that dmPFC neurons process contextual information during consolidation and retrieval, modulate discrimination between aversive and neutral contexts (Lingg et al., 2026; Rozeske et al., 2015, 2018), and that GR signaling in the dorsal dentate gyrus (dDG) and the lateral amygdala modulates engram allocation (Lesuis et al., 2021, 2025). This raises the possibility that post-training MIF infusion in the dmPFC could bias animals toward generalized freezing during retrieval by interfering with engram allocation during consolidation. Additionally, because remote contextual fear memories increasingly recruit mPFC networks during systems consolidation (Euston et al., 2012; Frankland & Bontempi, 2005; Matos et al., 2019), and because PrL and IL circuits maintain memory precision versus generalized fear responses over time (Pollack et al., 2018), we examined whether blocking dmPFC GRs after training alters remote fear memory strength and specificity. We tested separate groups of trained rats 2 or 14 days after CFC, exposing them sequentially to a novel context and then to the original training context, followed by perfusion and c-Fos immunohistochemistry (**Figure 2A**).

**Figure 2.**
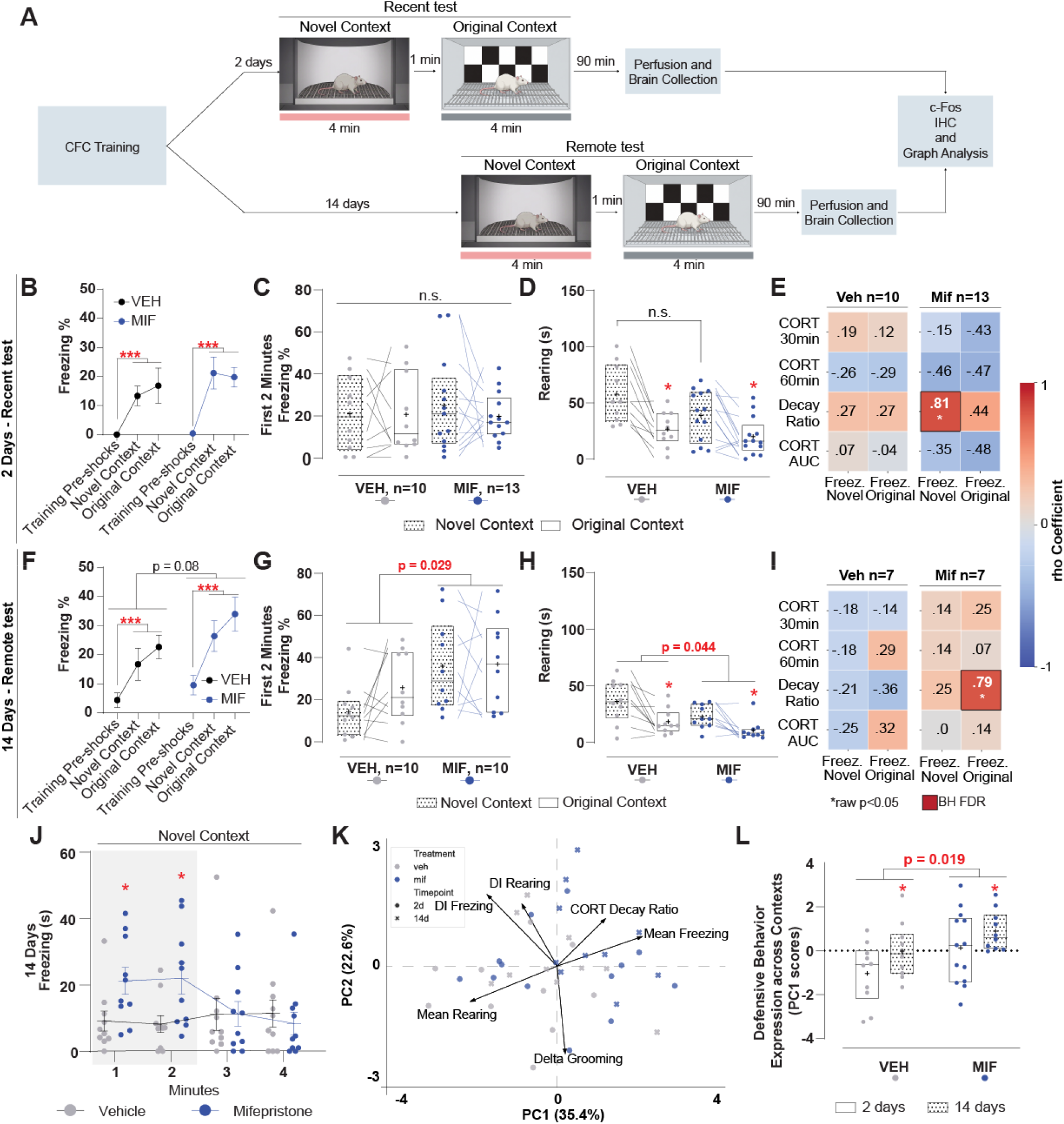
dmPFC GR blockade during consolidation spares recent but alters remote fear memory(A) Schematic of experimental design during the recent and remote memory tests. Two or 14 days after training, separate cohorts of rats were exposed to a novel context chamber for 4 min, followed by a 1-min interval and then by another 4-min exposure to the original training context. Freezing in response to the chambers was quantified during the whole test. Ninety minutes after the end of the test session, brains were collected for downstream immunohistochemistry and c-Fos graph analyses. **(B)** Mean freezing percentages ± SEM. during the pre-shock period of training, during exposure to the novel context, and during re-exposure to the training context at the 2 days test are shown for vehicle (VEH) and mifepristone (MIF) groups. Mixed ANOVA: Context F(2,42) = 18.02, p < 0.001, η²_G_ = 0.31; Treatment F(1,21) = 0.55, p = 0.47, η²_G_ = 0.01; Context*Treatment F(2,42) = 0.57, p = 0.57, η²_G_ = 0.01). ***p-value < 0.001 when compared to Training pre-shocks interval. **(C)** Median freezing percentages ± IQR in the novel and training contexts at the 2 days test. Datapoints from both groups (VEH and MIF) are shown in different colors (gray and blue respectively). Lines connect the different repeated measures values for individual animals, and boxplots showing median, interquartile intervals. The plus sign reflects the mean for each subset of data. Mixed ANOVA: Context F(1,21) = 0.17, p = 0.69, η²_G_ = 0.00; Treatment F(1,21) = 0.52, p = 0.48, η²_G_ = 0.02; Context*Treatment F(1,21) = 0.67, p = 0.42, η²_G_ = 0.01. **(D)** Median rearing duration (s) ± IQR in the novel and training contexts at the 2 days test, plotted as in C. Linear Mixed Model: Context F(1,21) = 27.74, p < 0.001; Treatment F(1,21) = 2.79, p = 0.11; Context*Treatment F(1,21) = 2.84, p = 0.11. Bracket represents Treatment comparison, *p-value < 0.001 when contrasting Contexts. **(E)** Heatmaps showing Spearman correlation coefficients (rho) between freezing measures (freezing during the novel and training contexts at test) and CORT parameters (30 min, 60 min, decay ratio, and AUC) for VEH (left) and MIF (right) groups. Color scale indicates direction and magnitude of correlations, with asterisks denoting coefficients that reach statistical significance and bold square borders indicating significant correlations that remain significant after Benjamini–Hochberg false discovery rate correction. **(F)** Mean freezing percentages ± SEM. during the pre-shock period of training, during exposure to the novel context, and during re-exposure to the training context at the 14 days test are shown for vehicle (VEH, black) and mifepristone (MIF, blue) groups. Mixed ANOVA: Context F(2,36) = 22.28, p < 0.001, η²_G_ = 0.39; Treatment F(1,18) = 3.45, p = 0.08, η²_G_ = 0.02; Context*Treatment F(2,36) = 0.74, p = 0.49, η²_G_ = 0.02. ***p-value < 0.001 when compared to Training pre-shocks interval. **(G)** Median freezing percentages ± IQR during the first 2 min for the novel and training contexts at 14 days. Mixed ANOVA: Context F(1,18) = 1.69, p = 0.21, η²_G_ = 0.03; Treatment F(1,18) = 5.61, p = 0.03, η²_G_ = 0.18; Context*Treatment F(1,18) = 1.20, p = 0.29, η²_G_ = 0.02. **(H)** Median rearing duration (s) ± IQR in the novel and training contexts at the 14 days test, plotted as in C. General Mixed Model: Context F(1,18) = 22.50, p < 0.001; Treatment F(1,18) = 4.71, p = 0.044; Context*Treatment F(1,18) = 0.86, p = 0.37. Bracket represents Treatment comparison, *p-value < 0.001 when contrasting Contexts. **(I)** Heatmaps with Spearman correlation coefficients (rho) for the remote timepoint, plotted as in (E). n = 7 per group. **(J)** Mean freezing duration analyzed minute by minute in the novel context at the 14 days test. Linear Mixed Model: Minute F(3,54) = 2.27, p = 0.09; Treatment F(1,18) = 1.59, p = 0.22; Minute*Treatment F(3,54) = 5.59, p = 0.002. *p-value < 0.05 significant interaction coefficients from Helmert contrasts comparing the treatment difference at each minute with the average treatment difference across subsequent minutes. **(K)** Principal component analysis (PCA) biplot showing individual animals and loadings for mean freezing, mean rearing, change in grooming, freezing and rearing discrimination indices (DIs), and CORT decay ratio. Color distinguishes treatment groups; circles and crosses distinguish recent and remote time points, respectively. The first and second principal components (PC1 and PC2) explain 35.4% and 22.6% of the variance, respectively. **(L)** Mean PC1 scores, labeled “Defensive Behavior Expression Across Contexts,” by treatment and time point. Open and patterned boxes indicate recent and remote tests, respectively. Linear Model: Treatment F(1,39) = 5.98, p = 0.019; Timepoint F(1,39) = 4.39, p = 0.043; Treatment*Timepoint F(1,39) = 0.09, p = 0.77. Bracket represents Treatment comparison, *p-value < 0.05 when contrasting Timepoints.

We quantified freezing, rearing, grooming and sniffing bouts and, at both timepoints, VEH and MIF-treated rats showed a higher proportion of freezing bouts and a lower proportion of rearing bouts in the original contexts than in the novel context (**Figure S1B**). At recent retrieval, behavioral category and context showed a significant association, with no evidence that this association differed between treatments. At remote retrieval, the context-behavior association differed between treatments, although both groups retained significant context-dependent differences in behavioral composition (both p<0.001). Grooming and sniffing accounted for relatively few bouts across conditions. Sniffing showed a significant context association only in MIF-treated rats at remote retrieval.

At the 2-days timepoint, for both VEH-(n = 10) and MIF-treated animals (n = 13), freezing increased markedly from the pre-shock phase of training to both recent test contexts, indicating successful acquisition and subsequent expression of contextual fear memory (**Figure 2B**). With relatively low levels of freezing during the tests, animals in both treatment groups exhibited similar freezing levels in the original context and novel contexts during the first 2 minutes, contrary to what would be expected for this footshock intensity at this time point (**Figure 2C**). Minute-by-minute freezing profiles across the 4-min exposures likewise did not reveal a clear treatment-dependent impairment in either the novel or training context (**Supplementary Figure 1C**). Both groups showed longer rearing duration in the novel context than in the original context, with no significant treatment effect or interaction between treatment and context (**Figure 2D**). Grooming duration showed no significant effects of treatment, context or their interaction (**Supplementary Figure 1D**). Correlation analyses identified a positive association between CORT Decay Ratio and novel-context freezing in MIF-treated rats that survived Benjamini-Hochberg correction. VEH-treated rats showed no significant associations between CORT measures and retrieval freezing (**Figure 2E**). Neither group showed significant correlations between CORT measures and rearing or grooming duration (**Supplementary Figure 1E**). These results suggest that post-training MIF treatment in the dmPFC does not clearly interfere with recent fear memory, but may alter the coupling between endocrine recovery after training and the later expression of context-specific fear.

At the 14-days timepoint, both VEH (n = 10) and MIF-treated rats (n = 10) showed higher freezing during the remote test than during the pre-shock time interval of the training session, indicating successful conditioning and persistent fear memory expression at this later timepoint (**Figure 2F**). During the first 2 minutes of the remote test, MIF-treated rats showed higher freezing levels than the VEH group in both the novel and original contexts (**Figure 2G**), suggesting that dmPFC GR blockade has its strongest behavioral effect on fear expression immediately upon context exposure at the remote timepoint. Minute-by-minute analyses revealed a Minute x Treatment interaction, with significant Helmert contrasts indicating greater MIF-VEH differences during the first 2 minutes than during subsequent exposure (**Figure 2J**). The full-session analysis also identified a Minute x Context x Treatment interaction, demonstrating that treatment-dependent freezing trajectories differed between contexts (**Supplementary Figure 1F**), suggesting increased fear generalization during initial exposure to a novel context. As at the 2-day timepoint, both groups showed longer rearing duration in the novel context than in the original context. Unlike at 2 days, MIF-treated rats showed shorter rearing duration overall than VEH-treated rats, without a significant Context x Treatment interaction (**Figure 2H**). Grooming duration showed no significant effect of Treatment, Context, or their interaction (**Supplementary Figure 1G**) Correlation analyses at the remote time point revealed a positive association between CORT Decay Ratio and freezing in the original context only for MIF-treated rats that survived Benjamini-Hochberg correction (**Figure 2I**). In VEH-treated rats, CORT AUC correlated negatively with grooming duration in the original context, although this association did not survive correction. No other correlations reached significance in either group (**Supplementary Figure 1H**).

To integrate behavioral and endocrine measures across retrieval timepoints, we performed principal component analysis (PCA) using mean freezing, mean rearing, changes in grooming, freezing and rearing discrimination indices, and CORT Decay Ration (**Figure 2K**). The first principal component (PC1) explained 35.4% of the variance and captured variation in overall defensive behavior, its contextual allocation and post-training CORT recovery. Higher PC1 scores reflected greater mean freezing, reduced mean rearing, lower freezing and rearing discrimination indices, and a higher CORT Decay Ratio. Only PC1 exceeded the permutation-derived retention threshold (**Supplementary Figure 1I**). MIF-treated rats showed higher PC1 scores overall than VEH-treated rats, and remote retrieval yielded higher scores than recent retrieval, with no significant Treatment x Timepoint interaction (**Figure 2L**). Together, these findings linked post-training GR blockade in the dmPFC promotes greater defensive responses that generalize across contexts at an early remote timepoint. We next asked whether these behavioral differences are mirrored by changes in functional co-activation among fear memory related brain regions.

### Post-training dmPFC GR blockade biases systems consolidation toward a denser, more interconnected retrieval network

Because c-Fos functional coactivity networks can provide a systems-level readout of how contextual fear memories engrams reorganize over time (Dos Santos Corrêa et al., 2022; Wheeler et al., 2013), and because glucocorticoid signaling modulates both synaptic and systems consolidation of fear memories (dos Santos Corrêa et al., 2021; Dos Santos Corrêa et al., 2019, 2022; Schwabe et al., 2022), we asked whether post-training dmPFC GR blockade changes how hippocampal, amygdalar, and cortical regions engage during recent versus remote retrieval. These regions support contextual fear memory retrieval and time dependent fear generalization (Cho et al., 2017; Frankland et al., 2004; Tonegawa et al., 2018; Tulogdi et al., 2012), and c Fos quantification can reveal their differential engagement during memory expression (Roy et al., 2022). We reasoned that MIF-induced changes in the CORT Decay Ratio during consolidation could extend beyond behavior to alter the broader systems-level recruitment of these regions across the retrieval time points. To test this, we quantified c Fos expression in hippocampal subregions (dDG, vDG, dCA3, vCA3, dCA1, vCA1), amygdalar nuclei (BLA, CeA), and cortical regions (ACC, PrL, aIC, aRSC) after retrieval, normalizing counts to home cage rats to compare VEH and MIF treated animals (**Figure 3A,B; Supplementary Figure 2A**). Whole-network mean correlation was highest in MIF-treated rats at 14 days compared to all other groups (**Figure 3C**).

**Figure 3:**
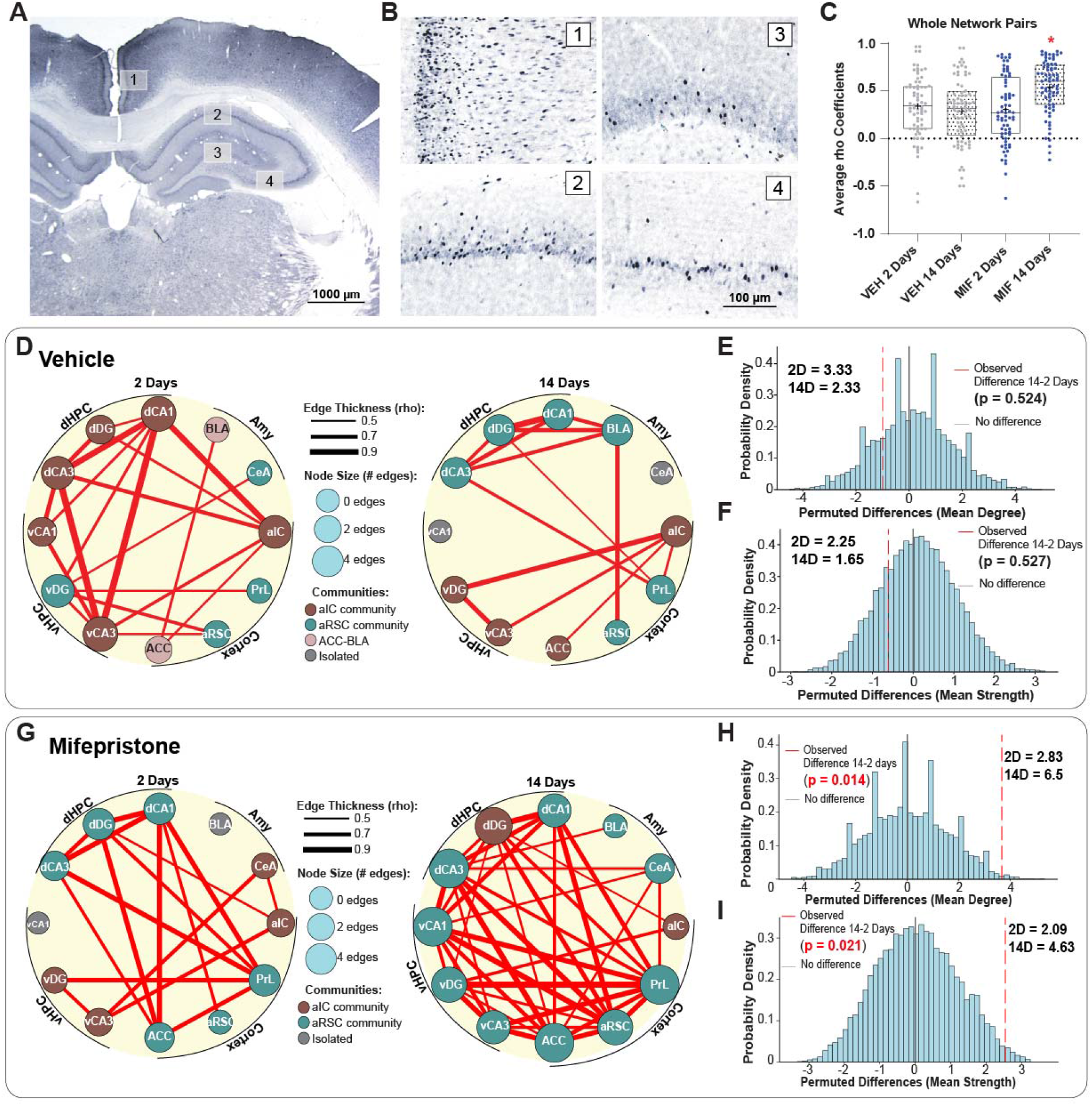
Post-training dmPFC GR blockade biases systems consolidation toward a denser, more interconnected retrieval network. **(A)** Representative coronal section illustrating regions sampled for c-Fos quantification. Numbers indicate sampled areas within dHPC, vHPC, Amy, and cortical regions on a DAB-stained slice. Scale bar, 1000µm. **(B)** Representative high-magnification images of c-Fos immunolabeling from the numbered regions in (A). These were used for cell counting. (1) aRSC, (2) dCA1, (3) dDG, (4) dCA3. Scale bars, 100µm **(C)** Median rho coefficient from all pairwise connections among the 12 sampled regions. (1-way ANOVA with bootstrap F[3,158] = 9.28, p < 0.001). *p-value < 0.05 compared to other groups. **(D)** Community structure and node degree organization of vehicle networks. Circular graphs depict thresholded co-activation networks at 2 days (left) and 14 days (right), where node size reflects the number of edges and edge thickness corresponds to rho magnitude. Nodes are colored by community membership. **(E-F)** Permutation-based comparison of global graph metrics between 2-day and 14-day vehicle networks. Histograms show null distributions of permuted differences for (E) mean degree per node, (F) mean strength per node. Observed differences are indicated by vertical red lines; gray vertical lines mark zero (no difference). Associated permutation p values and metric values for each time point are reported in the panels. **(G)** Community structure and hub organization of mifepristone networks, plotted as in D. **(H-I)** Permutation-based comparison of global graph metrics between 2-day and 14-day mifepristone networks, plotted as in E-F. Abbreviations: dHPC: dorsal hippocampus, vHPC: ventral hippocampus, Amy: amygdala, dDG: dorsal dentate gyrus, dCA3: dorsal CA3, dCA1: dorsal CA1, vDG: ventral dentate gyrus, vCA3: ventral CA3, vCA1: ventral CA1, BLA: basolateral amygdala, CeA: central amygdala, ACC: anterior cingulate cortex, PrL: prelimbic cortex, aIC: anterior insular cortex, aRSC: anterior retrosplenial cortex.

Building functional connectivity graphs requires binarizing correlation matrices with a fixed threshold; rather than the commonly used but arbitrary significance-based threshold method (Terstege & Epp, 2023), we calibrated this value empirically. We generated a permutation-based null distribution by randomly reassigning animals to four surrogate groups matched to the original sample sizes, then computed correlation matrices for each permutation, and concatenated all coefficients into a single null vector, which was plotted as a histogram next to the real data distribution (**Supplementary Figure 2B– C**). Real correlations were systematically shifted toward higher values than the null distribution, with the 70th percentile of the null distribution falling in the upper tail of the real distribution (**Supplementary Figure 2B–C**). Bootstrap resampling of mean real–null differences (**Supplementary Figure 2D**), together with Kolmogorov–Smirnov tests across permutations (**Supplementary Figure 2E**), confirmed that the real distribution was systematically and reliably shifted above the null distribution, supporting an empirically calibrated threshold rather than an arbitrary significance-based cutoff. Hence, we used the 70th percentile of this null distribution to build the binary identity matrices for all subsequent graph analysis. This approach applied the same edge threshold to all groups and based that threshold on the correlation values expected after randomly shuffling treatment and timepoint labels.

At 2 days, the VEH network seen in the functional connectivity matrix was characterized by prominent coordination among dHPC and vHPC nodes together with selected amygdalar and cortical regions (**Supplementary Figure 3A**). By 14 days, several pairwise relationships were redistributed (**Supplementary Figure 3B**), including notable changes in dCA3-vCA3, dCA1-vCA3, dDG-dCA1, and dDG-BLA coupling (**Supplementary Figure 3C**). Graph visualization supported this temporal reorganization for the VEH-treated rats, revealing changes in community structure and node centrality between the two timepoints (**Figure 3D; Supplementary Figure 3 D-E**). Community structure was identified using the Louvain algorithm on weighted adjacency matrices, which maximizes modularity to detect densely interconnected node groups. Excluding isolated nodes, Louvain identified three communities at 2 days (a larger hippocampus-aIC community, a medium vDG–CeA–PrL–RSC community, and a smaller BLA–ACC community) and two communities at 14 days (dHPC-BLA-PrL-RSC and vHPC-aIC-ACC) (**Figure 3D**). To test whether these topological differences exceeded chance, we compared each graph metric for observed networks with the null hypothesis of no group difference without assuming a theoretical distribution for the test statistics (Simpson et al., 2013). Despite the visible reorganization, mean degree (**Figure 3E**), mean strength (**Figure 3F**), average clustering (**Supplementary Figure 3F**), global efficiency (**Supplementary Figure 3G**), and modularity (**Supplementary Figure 3H**) all fell within their corresponding null distributions, indicating no global differences between the 2-day and 14-day VEH networks. This suggests that the time-dependent transition was expressed more through redistribution of connections and regional roles rather than a large-scale gain or loss of overall network integration. Together, these findings indicate that in VEH rats, 14 days retrieval is accompanied by reorganization of c-Fos co-activation without weakening or strengthening of the network, suggesting that under intact glucocorticoid signaling, the fear-memory network preserves overall functional coactivity while changing hippocampal-amygdalar-cortical interactions that support retrieval.

For MIF-treated rats, pairwise c-Fos correlation matrices showed robust interregional co-activation at both time points, but with a marked strengthening and redistribution of connectivity at 14 days, including broader positive coupling across hippocampal, cortical and amygdalar regions (**Supplementary Figure 4A-B**), and prominent ACC-BLA and ACC-vCA3 changes in the difference matrix (**Supplementary Figure 4C**). Graph visualization made this shift clear: the 2-day network showed a restricted set of strong links, whereas the 14-day network was denser and more integrated, with larger node sizes and more extensive cross-community connectivity (**Figure 3G; Supplementary Figure 4D-E**). Louvain clustering identified a larger dHPC-cortex and smaller vHPC-aIC-CeA community at 2 days, suggesting weaker intra-hippocampal but stronger hippocampal-cortical integration than VEH at the same timepoint, whereas at 14 days, with most regions collapsed into a single community (**Figure 3G**). Unlike VEH, this reorganization was accompanied by significant increases in mean degree (**Figure 3H**) and mean strength (**Figure 3I**) from 2 to 14 days, indicating a denser network with stronger surviving edges. By contrast, average clustering, global efficiency, and modularity did not significantly differ across time (**Supplementary Figure 4F-H**), suggesting that MIF’s main effect on systems consolidation was to enhance overall interregional coactivity rather than to fundamentally change large-scale network organization. Together, these findings show that MIF biases systems consolidation toward a stronger, more broadly interconnected 14-day retrieval network, consistent with a role in time-dependent fear generalization and susceptibility to PTSD (Dirven et al., 2026). Given this shift toward denser, less segregated coactivity, we next asked whether specific nodes disproportionately drive this reorganization by classifying regions into connector or provincial hub roles and by comparing coactivity in DMN-like versus SN-like subnetworks.

### Post-training GR blockade shifts DMN-like and Salience Network Contributions

To determine which subdivisions of the network were most affected by treatment, we quantified the average rho coefficient of pairwise c-Fos correlations for the whole network and for 4 subnetworks: within hippocampus (within HPC), hippocampus-cortex (HPC-CTX), hippocampus-amygdala (HPC-Amy), and amygdala-cortex (Amy-CTX) (**Figure 4A**). Subnetwork analyses revealed distinct temporal and treatment-dependent effects. Within-HPC coefficients were higher at 2 days than 14 days for VEH rats but higher at 14 days than 2 days in MIF rats, indicating that post-training dmPFC GR blockade reverses the temporal engagement of intra-hippocampal subregions during retrieval (**Figure 4B**). HPC-CTX coefficients were higher in MIF rats at 14 days than in VEH rats at either time point and higher in MIF rats at 2 days than in VEH rats at 14 days (**Figure 4C**). HPC-Amy coefficients increased from 2 to 14 days similarly in both treatments, indicating this connection is treatment-independent (**Figure 4D**), and Amy-CTX coefficients did not differ across any of the four groups (**Figure 4E**).

**Figure 4.**
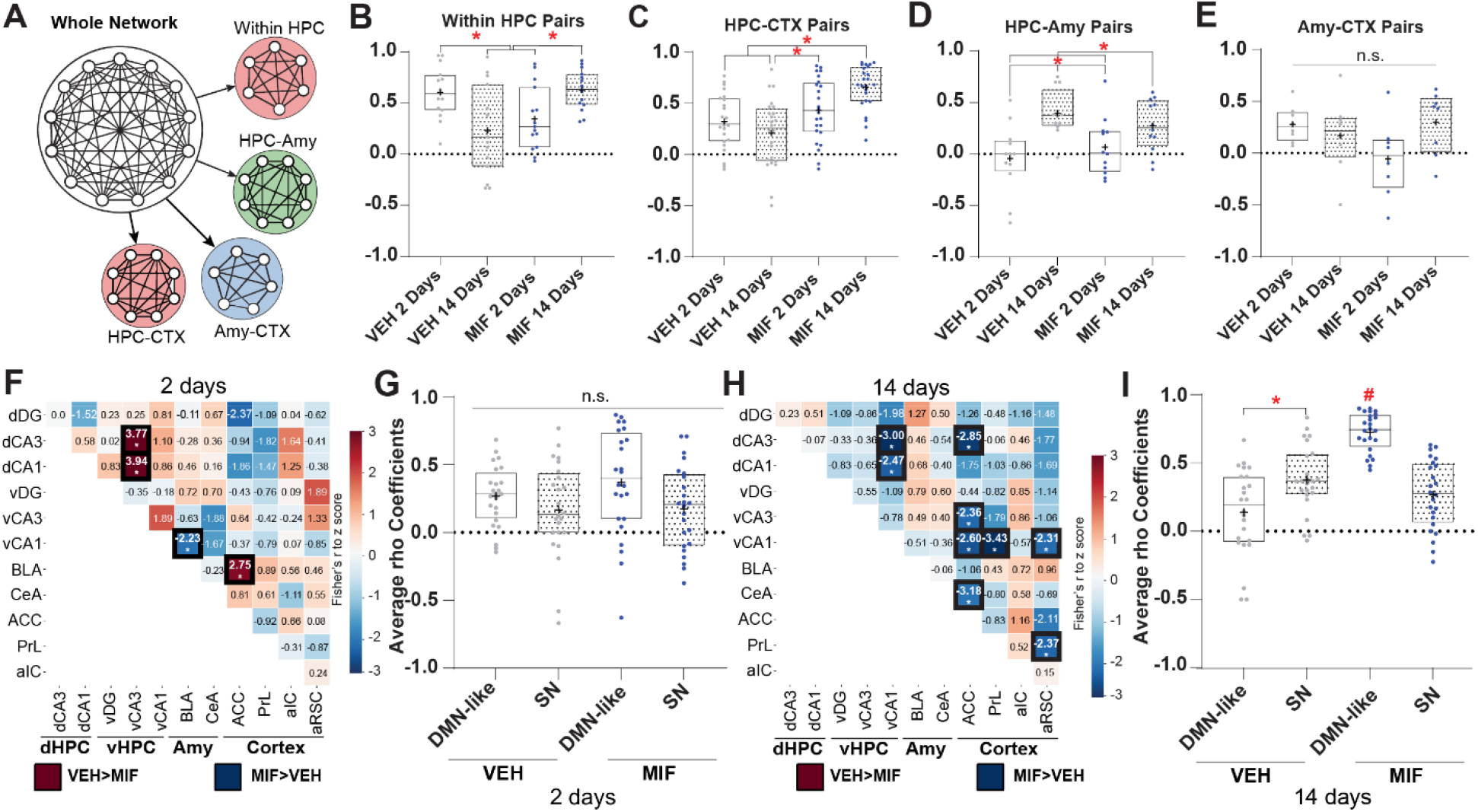
Post-training GR blockade shifts DMN-like and Salience Network Contributions. **(A)** Schematic of subnetwork definitions used for edge-wise analyses. We selected four subnetworks from the main graph (arrows). Within HPC pairs are all the pairwise connections between dorsal and ventral hippocampal subregions (dHPC×vHPC). HPC-Amy pairs are all the pairwise connections between hippocampal subregions and amygdala nuclei. HPC-CTX and Amy-CTX follow the same logic described previously, but between the referenced groups. **(B-E)** Boxplots show Spearman rho coefficients for individual c-Fos co-activation edges in vehicle-(VEH) and mifepristone-treated (MIF) rats tested at 2 days and 14 days. In each panel, dots represent individual edges, boxplots summarize group medians and interquartile ranges, the plus sign indicates the mean coefficient value; the dashed line denotes zero correlation. **(B)** Median rho coefficient from all pairwise connections among the edges between hippocampal subregions. (1-way ANOVA with bootstrap F[3,30.5] = 3.97, p = 0.022). *p-value < 0.05 compared to other groups as indicated by brackets. **(C)** Median rho coefficient from all pairwise connections among the edges between hippocampal subregions and cortical regions. (1-way ANOVA with bootstrap F[3,54.4] = 6.63, p = 0.005). *p-value < 0.05 compared to other groups as indicated by brackets. **(D)** Median rho coefficient from all pairwise connections among the edges between hippocampus subregions and amygdala nuclei. (1-way ANOVA with bootstrap F[3,24.3] = 5.56, p = 0.020). *p-value < 0.05 compared to other groups as indicated by brackets. **(E)** Median rho coefficient from all pairwise connections among the edges between amygdala nuclei and cortical regions. (1-way ANOVA with bootstrap, F[3,14.7] = 1.67, p = 0.216). *p-value < 0.05 compared to other groups as indicated by brackets. **(F)** Fisher *z*-score matrix showing pairwise differences in c-Fos co-activation (Spearman correlation) between mifepristone (MIF) and vehicle (VEH) groups at the 2 days retrieval test. Cells show the z-score for the VEH - MIF difference in each region pair; boxed values with asterisks indicate p < 0.05 (Fisher *r*-to-*z* transformation with two-tailed test, validated by permutation testing, n = 1,000 permutations). Higher z values (red) mean VEH has higher coefficient compared to MIF, whereas lower z values (blue) mean MIF has higher coefficient compared to VEH. **(G)** Average Spearman rho coefficients for edges within the default-mode-like (DMN-like) and salience network (SN) subnetworks in VEH and MIF groups at 2 days. Each dot represents one edge; box plots show median and interquartile range with “+” marking the mean. Bootstrapped one-way ANOVA: F(3,49.5)=1.85; p=0.15. n.s. denotes not significant. **(H)** Fisher z-score matrix of VEH - MIF differences in c-Fos co-activation 14 days after contextual fear conditioning, plotted as in (F). Boxed values with asterisks indicate p < 0.05. **(I)** Average Spearman rho coefficients for DMN-like and SN subnetwork edges in VEH and MIF groups at 14 days, plotted as in (G). Asterisks (red) denote significant pairwise differences between subnetworks and treatments. Bootstrapped one-way ANOVA: F(3,92)=31.26; p<0.001. *<0.05 in Bonferroni post hoc test compared to VEH SN. #<0.05 in Bonferroni post hoc test compared to all other groups. Abbreviations: dHPC: dorsal hippocampus, vHPC: ventral hippocampus, Amy: amygdala, dDG: dorsal dentate gyrus, dCA3: dorsal CA3, dCA1: dorsal CA1, vDG: ventral dentate gyrus, vCA3: ventral CA3, vCA1: ventral CA1, BLA: basolateral amygdala, CeA: central amygdala, ACC: anterior cingulate cortex, PrL: prelimbic cortex, aIC: anterior insular cortex, aRSC: anterior retrosplenial cortex.

Because global network metrics do not reveal which specific regions drive network reorganization, we next examined pairwise co-activation differences and node-level topological roles to determine how post-training dmPFC GR blockade reshaped region-specific contributions to the retrieval network, with a focus on hub distribution across DMN-like and SN subnetworks at each time point (Dirven et al., 2026). As main “seed” regions to define these subnetworks, we used the aRSC and ACC for the DMN-like (Lu et al., 2012; Nghiem et al., 2024) and the aIC for the SN (Zerbi et al., 2019), excluding conflicting edges (i.e. aIC-aRSC). At 2 days, Fisher z-transformed comparison of the MIF and VEH co-activation matrices revealed four significant pairwise differences after permutation testing: VEH increased dCA3-vCA3, dCA1-vCA3, and BLA-ACC coupling relative to MIF, while decreasing vCA1-BLA coupling (**Figure 4F**). Hub classification based on “within-module degree” and “participation coefficient” showed that in VEH animals, dCA3 and dCA1 acted as provincial hubs and vDG as a connector hub, whereas in MIF animals’ hub status expanded to include dCA3, dCA1, vCA3, and ACC as provincial hubs and PrL and dDG as connector hubs (**Supplementary Figure 5A**), indicating that hub regions spanned both DMN-like and SN nodes in both groups at this time point. Despite this broad hub distribution, average rho coefficients did not differ between DMN-like and SN subnetworks in either VEH or MIF groups at 2 days (**Figure 4G**), suggesting that MIF had not yet produced a detectable shift in the relative dominance of these two subnetworks at this time point. Together, these pairwise, hub-level, and subnetwork-level results indicate that at 2 days, post-training dmPFC GR blockade does not elicit a detectable shift in DMN-like versus SN subnetwork dominance.

By 14 days, the difference matrix showed a larger reorganization, with nine significant pairwise differences between MIF and VEH after permutation testing. MIF increased coupling within several hippocampal and hippocampal-cortical pairs, including dCA3-vCA1, dCA1-vCA1, dCA3-ACC, vCA1-ACC, vCA1-PrL, CeA-ACC, vCA3-ACC, vCA1-aRSC, and PrL-aRSC (**Figure 4H**). Hub classification at 14 days showed that VEH animals retained dCA3, dDG, and BLA as provincial hubs and aIC as a connector hub, while MIF animals shifted hub status predominantly to cortical regions with ACC, PrL, and aRSC acting as provincial hubs alongside residual hippocampal hub involvement from dCA3 and vCA1 (**Supplementary Figure 5B**), suggesting that DMN-like associated brain regions are more engaged than SN-like during remote retrieval for MIF-treated rats. This reorganization was accompanied by a significant subnetwork-level change: whereas VEH animals showed higher average connectivity in the SN than the DMN-like network, MIF animals showed the opposite pattern, with markedly elevated DMN-like connectivity exceeding SN connectivity (**Figure 4I**). Together, these results indicate that blocking post-training dmPFC GRs produced a progressive functional handoff from a SN weighted retrieval configuration in VEH-treated rats to a DMN-like dominated configuration at 14 days, establishing DMN-like hyperconnectivity as a defining feature of the remote retrieval network after memory consolidation under GR blockade. To determine whether these topological changes tracked behaviorally relevant outcomes, we next examined how region-specific c-Fos expression related to freezing behavior, fear generalization and CORT dynamics across timepoints and treatment groups.

### dmPFC GR blockade disrupts expected PrL’s association with fear memory consolidation

To determine whether post-training dmPFC GR blockade altered the recruitment and functional coupling of the same brain regions engaged during fear memory consolidation and retrieval, we compared c-Fos expression patterns and their correlations with behavioral and endocrine measures between VEH and MIF-treated rats across both timepoints. In VEH rats, retrieval was accompanied by region and time dependent differences in c Fos expression. At both timepoints, dCA3 and BLA showed higher expression relative to home cage animals. At 2 days, CeA and ACC also displayed higher c Fos levels, whereas vDG showed lower expression. At 14 days, dDG and vCA1 exhibited higher c Fos expression and no region showed lower expression relative to home cage controls. When comparing timepoints, dCA3, CeA, and ACC had lower expression at 14 days than at 2 days, whereas vDG and vCA1 increased their activity over time (**Figure 5A**). dCA1, vCA3, PrL, aIC, and aRSC did not differ significantly from home cage levels at either timepoint, nor did they differ between 2 and 14 days. We next computed correlation matrices between c Fos expression, freezing, and CORT measures for the VEH group and identified distinct patterns of association across timepoints. At 2 days, PrL c Fos expression was negatively correlated with several CORT measures (30 min, 60 min, decay ratio, and AUC) and positively correlated with post training freezing. At this timepoint, CeA c Fos was also inversely correlated with the CORT decay ratio. In contrast, at 14 days, c Fos in all vHPC subregions was negatively correlated with freezing in the novel context, and vCA1 expression was additionally negatively correlated with the generalization index (**Figure 5B**). A broadly similar, but not identical, pattern was observed after MIF treatment, suggesting that post training dmPFC GR blockade did not abolish network recruitment but altered how specific regions covaried with behavioral and CORT measures. When c Fos expression was compared to home cage animals, no region showed consistently higher or lower activity at both timepoints. At 2 days, dCA3, BLA, CeA, and ACC had higher c Fos expression than home cage animals, whereas vDG and vCA1 showed lower expression. At 14 days, no region in the MIF group differed significantly from home cage levels. In contrast, when comparing between timepoints, we found decreases over time in dCA3, BLA, and CeA and increases in vDG and vCA1. dDG, dCA1, vCA3, PrL, aIC, and aRSC showed no significant differences relative to home cage animals or between timepoints. Overall, the MIF treated group differed from the VEH group by showing a decrease over time in BLA activity instead of the ACC, effectively switching the region exhibiting reduced remote time activation (**Figure 5C**). Correlation matrices for the MIF group indicated that at 2 days there were no significant associations between c Fos expression and behavioral or CORT measures. At 14 days, however, we observed significant inverse correlations between freezing in the training context and c Fos expression in vDG, vCA3, and aIC. We also found positive correlations between the generalization index and PrL activity, and between the CORT decay ratio and dCA3 expression (**Figure 5D**). These findings suggest that MIF treatment may bias remote PrL activity toward generalized fear while shifting the contribution of the vHPC to remote memory retrieval.

**Figure 5.**
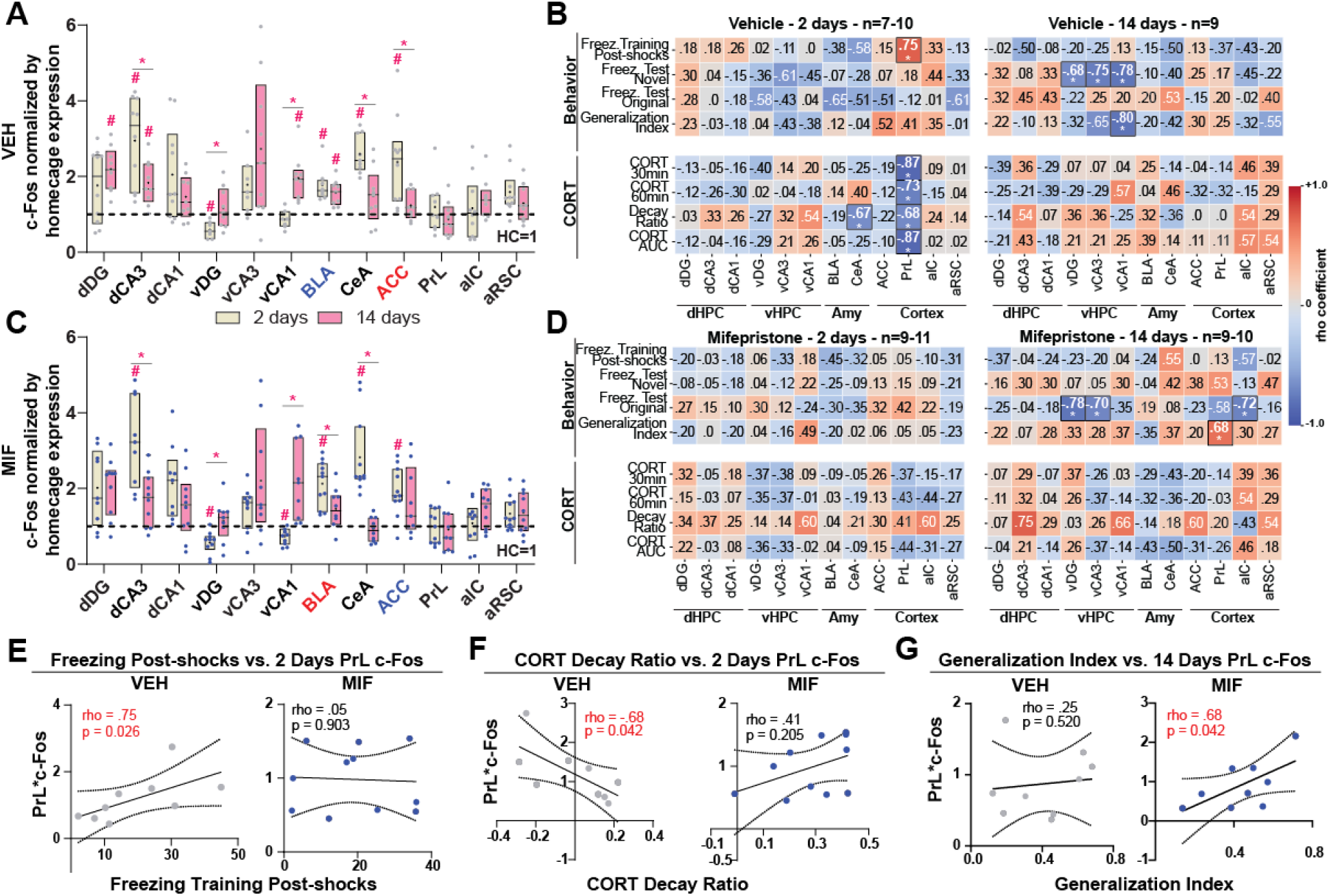
dmPFC GR blockade disrupts expected PrL’s association with fear memory consolidation. **(A)** Regional c-Fos expression in vehicle (VEH)-treated rats. Boxplots show c-Fos counts normalized to home-cage expression across hippocampal (HPC), amygdala (Amy), and cortical regions at 2 days (beige) and 14 days (pink) post-conditioning retrieval test. We used one-sample t-tests against the home-cage baseline (normalized to 1) to assess region-dependent changes, and two-sample t-tests to compare 2- and 14-day timepoints; we corrected all comparisons for multiple testing using FDR (# p<0.05 vs. home-cage baseline; * p<0.05 between timepoints). X-axis labels highlight regions with discrepant results between treatment groups: red indicates a region that reached significance in this treatment group but not in the other and blue indicates a region that reached significance in the other treatment group but not this one. **(B)** Heatmaps display rho coefficients between training and test freezing measures, generalization indices, CORT parameters (30 min, 60 min, decay ratio, AUC), and regional c-Fos expression at 2 days (left) and 14 days (right). Color scale indicates direction and magnitude of correlations, with asterisks denoting coefficients that reach statistical significance (raw and Holm-adjusted p values as indicated). **(C)** Regional c-Fos expression in mifepristone (MIF)-treated rats. Boxplots show c-Fos counts normalized to home-cage expression across HPC, Amy, and cortical regions at 2 days (beige) and 14 days (pink) post-conditioning retrieval test. We used one-sample t-tests against the home-cage baseline to assess region-dependent changes, and two-sample t-tests to compare timepoints, with FDR correction for multiple comparisons (# p<0.05 vs. home-cage; * p<0.05 between timepoints). Red and blue x-axis labels mark regions (BLA, ACC) that showed a significant effect in one treatment group but not the other. **(D)** Heatmaps with rho coefficients between the same behavioral, hormonal, and regional c-Fos variables as in (B) at 2 days (left) and 14 days (right) in MIF animals, with the same color coding and significance notation. **(E–G)** Targeted correlations linking CORT dynamics, contextual generalization, and PrL c-Fos activity in VEH (left) and MIF (right) groups. Each scatterplot shows data from individual animals with linear regression fits and 95% confidence intervals. Statistical results (rho and p values) are shown within each panel. **(E)** Relationship between post-shocks freezing in training session and PrL c-Fos expression at 2 days. **(F)** Relationship between CORT decay ratio and PrL c-Fos expression at 2 days. **(G)** Relationship between the generalization index at 14 days and PrL c-Fos expression during this timepoint. Abbreviations: dHPC: dorsal hippocampus, vHPC: ventral hippocampus, Amy: amygdala, dDG: dorsal dentate gyrus, dCA3: dorsal CA3, dCA1: dorsal CA1, vDG: ventral dentate gyrus, vCA3: ventral CA3, vCA1: ventral CA1, BLA: basolateral amygdala, CeA: central amygdala, ACC: anterior cingulate cortex, PrL: prelimbic cortex, aIC: anterior insular cortex, aRSC: anterior retrosplenial cortex.

To illustrate key group differences, we next plotted selected significant correlations and directly compared them to the corresponding correlations in the opposite treatment group (**Figure 5E-G**). In VEH, but not MIF treated rats, higher freezing during post-shocks phase of the training was associated with higher PrL c Fos expression at 2 days (**Figure 5E**). Similarly, only in the VEH group was a higher CORT Decay Ratio during the post-training consolidation window negatively correlated with PrL c-Fos expression at 2 days (**Figure 5F**). In contrast, in MIF treated rats, but not VEH, higher generalization at 14 days correlated positively with PrL c Fos expression (**Figure 5G**). Lastly, neither the 2-day recent nor the 14-day remote timepoint showed significant differences in c-Fos expression between VEH and MIF groups in any of the 12 brain regions examined (**Supplementary Figure 6A-B**). Together, these results suggest that post training dmPFC GR blockade decouples PrL activity from the acute stress response and fear consolidation signals present in VEH rats, while instead linking PrL engagement to generalized fear expression at the remote time point. This pattern may reflect disrupted GR-dependent regulation of CORT during the consolidation window and/or altered engram allocation within the dmPFC.

## DISCUSSION

In this study, we show that dmPFC GR signaling during the immediate post-learning window modulates the endocrine trajectory of the post-learning stress response and the eventual strength of contextual fear memory as it undergoes systems consolidation. Blocking GRs in the dmPFC during this window also shifted remote retrieval networks from an SN-like toward a DMN-like dominance, suggesting that dmPFC GR signaling sets the balance between specificity and generalization in contextual fear memory by shaping network-level trajectories as the memory matures.

Post-training GR blockade in the dmPFC altered the endocrine trajectory of the stress response, accelerating CORT decay between 30 and 60 minutes without changing total CORT output (**Figure 1E, Supplementary Figure 1A**), and this change coincided with a shift in fear memory expression that appeared only at the remote, not the recent, time point (**Figure 2C-D,G-H**). At 14 days, MIF-treated rats spent, on average, more time freezing than VEH-treated rats in both contexts, an effect driven specifically by higher freezing during the initial minutes of novel context exposure rather than increased freezing overall (**Figure 2J**). PCA integrated these behavioral changes with contextual discrimination and endocrine recovery revealing higher scores in MIF-treated rats on a component characterized by greater freezing, reduced rearing, lower discrimination indices, and faster CORT decay, implying that dmPFC GR blockade promotes generalized defensive behavior expression **(Figure 2K-L)**. This behavioral phenotype occurred alongside with a shift at the network level: VEH retrieval networks showed SN dominance at 14 days, whereas MIF networks shifted to a DMN-dominant configuration (**Figure 4I**), with hub status moving from hippocampal and amygdalar nodes toward cortical regions, including ACC, PrL, and aRSC – canonical DMN regions (**Figure 4B-E**). This reorganization was accompanied by a progressive densification of the MIF network between 2 and 14 days, reflected in significant increases in mean degree and mean strength **(Figure 3H,I)** that were absent in VEH networks over the same interval (**Figure 3E-F**), indicating that GR blockade produced a more globally interconnected retrieval network rather than a change confined to a small set of hub regions. Within this reorganized network, PrL activity showed a region-specific change: in VEH rats, PrL c-Fos expression during recent retrieval correlated negatively with CORT measures and positively with post-shock freezing; these associations did not reach significance in MF rats, in which remote PrL expression instead correlated positively with generalization (**Figure 5 B/E-G**). Together with its hub status in the remote MIF network, this shift identifies PrL as a candidate node linking the loss of GR-dependent CORT coupling during consolidation to the altered network dominance and increased freezing levels observed at remote retrieval.

The CORT decay ratio also correlated positively with training freezing selectively in MIF-treated rats. This faster CORT decay under MIF departs from a classical negative-feedback interpretation, since the mPFC ordinarily provides GR-dependent feedback inhibition onto the HPA axis via projections to the BNST (Herman et al., 2016; Radley, 2012; Ulrich-Lai & Herman, 2009), and blocking dmPFC GRs would be expected to impair, rather than accelerate, this feedback. No prior study has directly examined how dmPFC-specific GR blockade affects HPA axis response: a comprehensive review of mifepristone-HPA axis studies (Dalm et al., 2019) identified no prior investigation targeting the dmPFC specifically, highlighting that our data begin to address an open question. Two, not mutually exclusive, interpretations remain viable. First, dmPFC GR blockade may facilitate HPA axis negative feedback through a compensatory pathway involving other feedback sites, such as the hippocampus, that remain unblocked (Herman et al., 2016; Jacobson & Sapolsky, 1991). Second, GR blockade may shift the local GR/MR ratio toward MR dominance, as shown for the hippocampus (Ratka et al., 1989; Van Haarst et al., 1997), facilitating dmPFC-BNST communication; under this account, the faster CORT recovery observed after MIF treatment would track increased MR engagement in the dmPFC. Further experiments manipulating MR signaling directly in the dmPFC will be needed to test this possibility. The pattern we found parallels results reported by Van Haarst et al. (1997) for dHPC GR blockade (Van Haarst et al., 1997), and is consistent with a shift toward MR dominance facilitating negative feedback in that region. GR and MR receptors more broadly modulate the behavioral strategies animals adopt in goal-directed tasks (Oitzl & de Kloet, 1992), and a similar logic may extend to contextual fear generalization: stress is proposed to shift memory consolidation toward habit-based, less flexible learning at the expense of cognitively precise memory (Quaedflieg & Schwabe, 2018), a shift potentially driven by MR activation in the relative absence of functional GR signaling (Asano et al., 2025; Vogel et al., 2016). Following this logic, our finding that GR blockade in the dmPFC does not impair fear conditioning consolidation but instead facilitates fear expression at a remote time point is consistent with a GR/MR shift toward MR dominance favoring consolidation of a stronger memory trace. However, this interpretation does not establish that altered endocrine recovery mediates the subsequent memory effects. Recent work demonstrates that prefrontal regulation of endocrine responses and memory specificity can dissociate. Post-training inhibition of the rostral PrL projections to the anteroventral bed nucleus of the stria terminalis pathway promotes contextual generalization even when animals are adrenalectomized, indicating that a generalized fear memory trace may occur independently from CORT levels during memory consolidation (Lingg et al., 2026). Thus, the endocrine and behavioral consequences of dmPFC GR blockade may likewise reflect partly separable processes.

At the network level, acute stress reorganizes large-scale brain networks in a time-dependent manner, favoring an SN configuration involving the amygdala and dmPFC in the immediate aftermath of a stressor (Hermans et al., 2011), and shifting toward a DMN-like configuration during the delayed, corticosteroid-dependent phase (Schwabe et al., 2022). The “cortisol switch” framework holds that CORT orchestrates coping and adaptation via a receptor-mediated on/off mechanism in which MRs and GRs jointly coordinate defensive reactions across distributed stress circuitry, and that vulnerability arises when this balance or its temporal dynamics is disrupted (de Kloet & Joëls, 2023). Consistent with a role for this receptor balance in network organization, our findings suggest that dmPFC GR signaling during post-training consolidation influences the later balance between SN-like and DMN-like engagement at retrieval. This extends the broader view that glucocorticoid signaling regulates shifts between vigilance-oriented and integrative, consolidation-oriented network configurations (de Kloet & Joëls, 2023; Hermans et al., 2014; Schwabe, 2017). The combination of elevated freezing to a safe, novel context and DMN-like hyperconnectivity in the remote retrieval network parallels a core feature of PTSD and related anxiety disorders, in which patients experience aberrant fear responses to safe cues or contexts that merely resemble a traumatic one (Kaczkurkin et al., 2017; Lissek et al., 2005; Webler et al., 2021). Supporting the translational relevance of this framework, resting-state functional networks analogous to the human SN and DMN have recently been identified in rodents, indicating that these large-scale network configurations are conserved across species and can be probed directly through rodent models (Grandjean et al., 2020; Zerbi et al., 2015). In a complementary severe-stress model, susceptible mice showed increased SN-DMN coupling around and after stress exposure, greater intra-SN connectivity after stress, and heightened RSC activity relative to resilient mice (Dirven et al., 2026). Although this pattern differs from the DMN-like dominance we observed at remote memory retrieval, both studies associate adverse behavioral outcomes with altered organizations of salience and default-mode related circuitry. Our data identify post-training dmPFC GR signaling as one regulator of this later network balance engaged during memory retrieval, providing a candidate mechanism for investigating how early responses to aversive experience shape trauma-related memory overgeneralization.

Our analytical framework also strengthens the interpretation of these network-level effects. Rather than applying a conventional fixed correlation or significance threshold, we calibrated graph binarization against an empirical null distribution generated by label permutation and tested global metrics through repeated animal-level reassignment with complete graph reconstruction. This approach reduces dependence on arbitrary edge selection and evaluates whether the observed differences in network density and strength exceed variation expected from the sampled animals and group sizes. Although c-Fos coactivity remains a correlational, group-level estimate of functional organization, the convergence of data-driven thresholding and permutation-based inference supports the conclusion that MIF reorganized remote retrieval-network topology. Additionally, this framework provides a reproducible strategy for testing c-Fos coactivity-network hypotheses when sample sizes are necessarily modest and threshold-dependent graph metrics may otherwise be difficult to interpret.

Two, not mutually exclusive, alternative accounts merit consideration alongside this network-level framework. The first concerns engram allocation rather than network-level reorganization per se. Glucocorticoid signaling in the dDG and lateral amygdala biases engram allocation toward larger, less selective neuronal ensembles, a mechanism proposed to underlie stress-induced fear generalization (Lesuis et al., 2021, 2025). If a comparable process occurs in the dmPFC, GR blockade could bias the size or composition of the dmPFC engram allocated during consolidation, and the DMN-like hub shift we observe in ACC, PrL, and aRSC at 14 days would reflect the functional signature of this altered engram rather than a cause of altered fear specificity in its own right. The shift in PrL’s behavioral correlates we observe, from an association with CORT and post-shocks freezing during training in VEH rats to an association with the generalization index in MIF rats, is consistent with, though does not directly demonstrate, this possibility. Distinguishing an engram-allocation account from a network-reorganization account will require activity-dependent tagging approaches (Takehara-Nishiuchi, 2021), to resolve whether dmPFC GR blockade changes ensemble size or overlap independent of its downstream network consequences.

The second alternative concerns the quality of prefrontal-hippocampal communication during consolidation. GR activation in the ACC enhances stress-sensitive high-frequency network oscillations ex vivo (Donaire et al., 2025), raising the possibility that dmPFC GR blockade altered oscillatory coupling between the dmPFC and hippocampus during the immediate post-learning window, independent of its effects on CORT decay. Because hippocampal-cortical oscillatory synchrony is thought to support the transfer and stabilization of contextual information during memory consolidation (Maingret et al., 2016), a loss of GR-dependent oscillatory activity in the dmPFC could degrade the fidelity of information transferred from the hippocampus (Schmidt et al., 2019), producing a less context-specific memory trace independent of the SN-to-DMN shift described above. This account is compatible with our c-Fos findings, since disrupted oscillatory coupling during consolidation could plausibly manifest downstream as the altered hippocampal-cortical coactivity we observe in the MIF group at 14 days (Figure 3F). Direct electrophysiological recordings during the post-training window will be needed to test this possibility.

These mechanistic possibilities raise questions within the broader context of our findings on CORT and fear memory consolidation. Our own prior work showed that CORT does not modulate CFC consolidation when training involves a single 0.6 mA footshock (Bueno et al., 2017), but facilitates time-dependent fear generalization when administered after training with three footshocks of the same intensity (dos Santos Corrêa et al., 2021), and that these protocols engage distinct brain regions and produce different temporal dynamics of contextual specificity during remote retrieval (Dos Santos Corrêa et al., 2019, 2022). This suggests that CORT acts secondarily to other physiological processes triggered by emotional activation, such as noradrenergic release and endocannabinoid signaling, which may set the initial direction of engram allocation that CORT subsequently amplifies (Atucha et al., 2017; Corrêa et al., 2023; Zerbi et al., 2019). This framework situates our present finding that blocking dmPFC GRs altered a component of the network recruited during engram allocation for mild to moderate intensity training.

This raises the question of what role the dmPFC plays in contextual specificity, particularly during recent retrieval. While early lesion work assigned generalized remote fear retrieval specifically to ACC rather than PrL (Frankland et al., 2004), subsequent optogenetic studies show that PrL engram cells allocated during acquisition become behaviorally relevant specifically at remote, not recent, retrieval (Kitamura et al., 2017), and that PrL engagement in contextual specificity depends on training intensity (Matos et al., 2019). Together, these studies suggest that dmPFC involvement in remote retrieval extends beyond time-dependent fear generalization and depends on both the subregion and the learning conditions. Our findings extend this temporal distinction by showing that, at recent retrieval, post-training dmPFC GR blockade altered selected interregional c-Fos correlations despite no significant treatment effects on freezing or rearing (**Figures 2C-D and 4F**). These findings indicate that differences caused by our treatment in memory retrieval coactivation can occur without detectable changes in these behavioral outcomes.

In summary, our study reveals a previously unrecognized role for dmPFC GR signaling during memory consolidation in regulating the transition from an SN-like to a DMN-like retrieval network configuration, thereby limiting the emergence of fear responses to safe, novel contexts as the memory becomes remote. Fear overgeneralization is a defining feature of PTSD and related anxiety disorders, and advancing treatment strategies for these conditions requires a fundamental understanding of the neuroendocrine mechanisms that constrain fear memory generalization. Insights gained from our results can inform the development of models linking early post-trauma endocrine dysregulation to later memory generalization, ultimately supporting the design of interventions that target the post-learning consolidation window in trauma-related psychopathology.

## LIMITATIONS OF THE STUDY

This study has several limitations, though our design and analysis choices help offset each one. In terms of methods, most cannula placements targeted the PrL, with only partial coverage of the ACC. This prevented a direct comparison of GR-dependent contributions between these two dmPFC subregions. Nonetheless, histological verification confirmed consistent targeting of the dmPFC across nearly all animals, and our findings converge with prior work implicating PrL specifically in remote memory precision (Matos et al., 2019), strengthening the interpretability of the PrL-associated effects we report. Additionally, the c-Fos coactivation network analysis is only correlational, co-activation patterns and their associations with behavior and CORT dynamics indicate covariation across regions but do not establish directional or causal connectivity between nodes. Even so, because network structure was compared between a pharmacologically manipulated (MIF) and control (VEH) group, any topological differences can still be attributed to dmPFC GR post-training blockade itself, giving the network analysis causal support that purely observational c-Fos studies lack. Lastly, only male rats were used, so whether these findings extend to females, in whom HPA axis reactivity and GR signaling differs, is unknown. Restricting the design to males reduces variability introduced by estrous-cycle fluctuations in CORT and GR expression (Oyola & Handa, 2017), allowing us to isolate dmPFC GR-specific effects on consolidation with greater statistical and mechanistic clarity, providing a controlled foundation before extending the model to females.

More importantly, we did not directly test engram allocation (e.g., via activity-dependent tagging at acquisition), so the proposed link between post-training GR blockade and the allocated dmPFC fear engram remains inferential. This limitation is mitigated by the fact that our correlational findings (PrL recent c-Fos expression is inversely correlated with CORT secretion but remote expression is associated to generalized fear) are consistent with prior engram-based evidence (Matos et al., 2019; Rao-Ruiz et al., 2021), generating a specific, testable hypothesis for future tagging studies rather than relying on unsupported speculation. Moreover, overall freezing generalization did not differ significantly between VEH and MIF groups at either timepoint, indicating that the MIF-induced increase in freezing reflects heightened overall memory strength rather than a selective change in context discrimination. However, we assessed additional behaviors beyond freezing and used PCA to characterize the treatment effect across multiple behavioral measures. These analyses complement statistically robust elevation in freezing and the clear reorganization of SN-like/DMN-like network topology at 14 days provide convergent, independently significant evidence that dmPFC GR blockade meaningfully alters the remote memory trace. Finally, c-Fos was sampled at a single retrieval timepoint per animal (2 or 14 days), providing a static snapshot of network engagement rather than a within-subject trajectory. This cross-sectional design nonetheless enabled us to directly compare distinct cohorts at biologically meaningful recent and early remote consolidation time points.

## RESOURCE AVAILABILITY

### Lead contact

Further information and requests for resources should be directed to the lead contact, Raquel Vecchio Fornari.

### Materials availability

No new physical materials were generated by the computational analyses described here.

### Data and code availability

- Processed behavioral data (freezing percentages across training and retrieval sessions), plasma corticosterone (CORT) measurements, c-Fos cell counts across hippocampal, amygdalar, and cortical regions, processed pairwise Spearman correlation matrices, and graph-theoretic network metrics (node degree, strength, clustering, modularity, participation coefficient), along with JAMOVI (.omv) session files and full experiment metadata, have been deposited at Mendeley Data and are publicly available as of the date of publication at: Dos Santos Corrêa, Moisés; Fornari, Raquel (2026), “Data for ‘Prefrontal cortex glucocorticoid receptors during fear memory consolidation shift the balance between salience and default-mode networks at retrieval in rats,’” Mendeley Data, https://doi.org/10.17632/gvykybtvw7.
- Raw c-Fos immunohistochemistry images and raw behavioral video recordings cannot be deposited in a public repository due to file-size limitations. These datasets will be shared by the lead contact upon reasonable request.
- All original analysis code used in this study, including permutation-based threshold calibration, graph construction, hub classification, and default-mode-like/salience-network-like subnetwork analyses, has been deposited at GitHub and archived at Zenodo, and is publicly available as of the date of publication at: https://doi.org/10.5281/zenodo.22025170 (GitHub repository: https://github.com/violettiger/dmPFC_MIF). The computational workflow was implemented in Python using four analysis files: network_complete.py, permutation_analysis.py, subnetworks.py, and Behavioral_correlations.ipynb. The scripts read local tabular input files and export CSV, PNG, and EPS outputs.
- Any additional information required to reanalyze the data reported in this paper is available from the lead contact upon request.

## ACKNOWLEDGMENTS

We dedicate this manuscript to Iraci Kosby Corrêa, in memoriam. This work was supported by the São Paulo Research Foundation (FAPESP) grant #2017/03820–0 (RVF), FAPESP fellowship #2017/24012–9 (MdSC), FAPESP grant #2023/07864-2 (LVL), Alexander von Humboldt Foundation, Germany (fellowship to M.d.S.C.), and *Conselho Nacional de Desenvolvimento Científico e Tecnológico* (CNPQ) grant #406075/2021-2 (TLF). We also thank the work done by the technicians that manage the vivarium. The authors would like to thank the members of the Research group in Neurobiology of Learning and Memory at UFABC (MANAs) for scientific advice and discussion.

## AUTHOR CONTRIBUTIONS

Conceptualization, M.d.S.C., P.A.T and R.V.F.; formal analysis, M.d.S.C. with inputs from P.A.T, and R.V.F.; investigation, M.d.S.C., L.V.L., A.C.Q.d.S., J.C.C., W.T.B.L., A.C.C.S., T.L.F., and R.V.F.; resources, M.d.S.C, R.V.F, and T.L.F.; visualization, M.d.S.C., with inputs from all authors; writing – original draft, M.d.S.C., with inputs from all authors; and supervision, P.A.T, and R.V.F.

## DECLARATION OF GENERATIVE AI AND AI-ASSISTED TECHNOLOGIES IN THE WRITING PROCESS

During the preparation of this work the authors used Perplexity© in order to improve the readability and language of the manuscript and to improve the Python code used for data analysis. After using this tool/service, the authors reviewed and edited the content as needed and take full responsibility for the content of the published article.

## SUPPLEMENTAL INFORMATION

Document S1. Figures S1–S6

## EXPERIMENTAL MODEL AND STUDY PARTICIPANT DETAILS

Three-month-old male Wistar rats, obtained from Instituto Nacional de Farmacologia (INFAR-UNIFESP (total n = 52; 6 home-cage; weighing at least 270 g at time of surgery), were kept in controlled conditions of temperature (23 ± 2 °C) and 12-h light/darkness cycle (light phase starting at 7 a.m.) with food and water ad libitum. In the study, “n” refers to number of animals. The rats were randomly house four per cage (40 cm x 33cm x 18 cm) and acclimated to the vivarium at the Universidade Federal do ABC (UFABC), São Bernardo do Campo campus, for at least one week before the beginning of the experiments. Following surgery, the animals were housed individually (20 cm × 16 cm × 18 cm) during the initial recovery period and subsequently housed in pairs, two per cage, for the remainder of the experiment. Behavioral experiments were performed during the light phase of the cycle, between 10 a.m. and 3 p.m., corresponding to the nadir of the CORT circadian rhythm. All procedures were conducted according to the guidelines and standards of CONCEA - Conselho Nacional de Controle de Experimentação Animal (Brazilian Council of Animal Experimentation) and were previously approved by the Ethics Committee on Animal Use of the Federal University of ABC (CEUA-UFABC; protocol number 6932250621). Reporting in this manuscript follows ARRIVE guidelines 2.0.

## METHOD DETAILS

### Stereotaxic surgery

The rats underwent stereotaxic surgery for bilateral cannula implantation as previously described (de Paiva et al., 2021). As a pre-anesthetic, the animals received a subcutaneous injection of acepromazine (1 mg/kg) and, after 15 minutes, were anesthetized with an intraperitoneal solution of ketamine and xylazine in sterile saline (90 mg/kg and 9 mg/kg, respectively). The animals were positioned in a stereotaxic apparatus and, following skull exposure, stainless-steel guide cannulas (11 mm, 23 gauge) were implanted bilaterally into the dorsomedial prefrontal cortex (dmPFC) following the coordinates: 3.2 mm anterior to bregma, 0.7 mm lateral to the midline, and 2.0 mm ventral to the skull surface (based on Paxinos & Watson, 2007). The cannulas were secured to the skull using two anchoring screws and dental cement, and a stainless-steel stylet (11 mm long) was inserted into each cannula to prevent obstruction. At the end of the surgery, the animals received an injection of antisedan (0.5 mg/kg) as a sedative reversal agent, 3 mL of sterile saline for hydration, meloxicam (2 mg/kg) for anti-inflammatory purposes, enrofloxacin (0.1 mL) as an antibiotic, and rifamycin spray on the edges of the cement. Animals were placed in individual heated boxes for recovery from anesthesia, and 5 mL of dipyrone was added to their 200 mL water bottles to assist with pain management. Finally, the animals were returned to the vivarium in individual transparent cages that were placed side by side to reduce the perception of isolation. The minimum recovery period before behavioral testing was 7 days, during which animals were monitored for weight gain, suture condition, cannula placement, signs of infection, and locomotor activity. Signs of pain were resolved with supplemental analgesic administration (meloxicam, 1 mg/kg).

### Contextual fear conditioning task and drug infusion

Behavioral experiments were conducted in two identical automated fear-conditioning chambers (Med Associates, Inc., St. Albans, VT), equipped with a grid floor connected to an Aversive Stimulator (ENV-414S), sound stimulus generators and infra-red-light camera (VID-CAM-MONO-4 Fire Wire Video Camera), as well as to a computerized interface that enabled video recording, real-time analysis, and measurement of the rats’ freezing behavior, as previously described (Dos Santos Corrêa et al., 2019). Freezing behavior was analyzed using Video Freeze software (Version 1.12.0.0, Med Associates), which quantified the cumulative duration during which movement remained below a predefined threshold of 20 arbitrary units for at least 1 s. This duration was considered freezing behavior. The software also outputs the Average Motion Index, the mean frame-by-frame motion value (a.u.) across a given component, used as a continuous measure of general activity. The conditioning chambers (32 cm wide, 25 cm high, and 25 cm deep, VFC-008) had transparent polycarbonate front and top walls, while the rear and side walls were made of white acrylic. Each chamber was housed within a sound-attenuating enclosure (NIR-022SD; 63.5 cm × 35.5 cm × 76 cm) and illuminated by an LED light source (NIR-100; Med Associates) that emitted both visible white light (450–650 nm) and near-infrared light (940 nm).

Contextual fear conditioning (CFC) training and testing sessions were conducted as previously described (dos Santos Corrêa et al., 2021). Animals were trained in a fear-conditioning chamber (Original Context) characterized by 5-cubed white-on-black pattern at the back wall, an exhaust fan noise, white illumination, and a stainless-steel grid floor with parallel rods through which footshocks were delivered. The Original Context was cleaned with 10% ethanol before and after each session. Before training, animals were handled for 3 consecutive days to habituate them to the experimenter and all experimental procedures. On the training day, animals remained in a room next to the conditioning room for at least 1 hour before CFC. During training, rats were placed in the Original Context and, after 2 min, received three footshocks (0.6 mA, 1 s duration) delivered at 30 s inter-shock intervals. Animals were removed from the chamber 1 min after the last footshock, resulting in a total session duration of approximately 4 min.

Immediately after training, the animals received bilateral infusions of either mifepristone (MIF) or vehicle into the dmPFC. We allocated rats to treatment groups using a pseudo-random manner, stratified to balance baseline body weight and cannula placement across groups. The lead author was aware of group allocation whereas experimenters were blinded to the groups during CFC task, and blood collection. Microinfusions were performed using 30-gauge injection needles connected via polyethylene tubing to 5-μL Hamilton microsyringes. The injection needles extended 2 mm beyond the tips of the guide cannulae, and a volume of 0.5 μL was infused over a period of 60 s using an automated infusion pump. The injection needles were left in place for at least 20 s after the end of the infusion to prevent reflux into the cannulae. MIF (Sigma-Aldrich) was initially dissolved in 100% ethanol and subsequently diluted in sterile phosphate-buffered saline (PBS) to achieve a final ethanol concentration of 0.5%. The final concentration of MIF was 20 ng/μL, and 0.5 μL per hemisphere was used for microinfusion (total of 10ng of MIF per hemisphere). An equivalent volume of vehicle solution (0.5% ethanol in PBS) was used as the control treatment (VEH). Following the infusions, animals remained in their individual cages and were transported to a separate room (collection room), where tail blood samples were collected 30 and 60 min after training. After these procedures, each animal remained in its individual cage in the collection room for at least 1.5 h before being returned to its group cage and replaced on the vivarium racks. The home-cage group did not undergo surgery, experimenter handling, or CFC training. However, these animals underwent tail blood collection following the same schedule as the trained groups.

CFC testing was conducted 2 or 14 days after training and consisted of sequential exposures to the Novel Context and the Original Context, each lasting 4 min and separated by a 1-min interval. The Novel Context was characterized by a flexible white acrylic placed at the back of the chamber to create a semicircular configuration. The grid floor was composed of 20 interleaved stainless-steel rods of either 4.8 or 9.5 mm of diameter. In addition, the chamber light remained off, 90-dB white noise was continuously presented, and the chamber was cleaned with a 3% acetic acid solution before each session (dos Santos Corrêa et al., 2021). During exposure to the Novel Context, the lights in the experimental room remained off whereas during exposure to the Original Context, lights in the room were on. No footshocks were delivered during testing, and freezing behavior was automatically recorded throughout both sessions.

### Behavioral Annotation

We annotated rearing, grooming and sniffing offline from the video recordings of each retrieval session using BORIS (Behavioral Observation Research Interactive Software, v(XX)) (REF). We set up an ethogram in which each behavior was a mutually exclusive state event, and we logged when each bout started and ended. Video scoring was done at normal playback speed, and the observer was blinded to treatment and retention interval. We defined the behaviors as follows:

- Rearing: the animal raises both forepaws off the floor and stands upright on its hind limbs. We counted both supported rearing (forepaws against the chamber walls) and unsupported rearing (forepaws in the air).
- Grooming: the animal licks or strokes its forepaws, face, head or body, or scratches itself with a hind paw. This includes the full head-to-tail grooming sequence. A bout ended when grooming stopped for more than 1s.
- Sniffing: the animal moves its head, with the nose pointed at the floor, walls or air, and its whiskers move. It can happen as the animal walks or stands still, but not during rearing, grooming or freezing.

For each behavior we calculated total duration in seconds. We exported the time budgets from BORIS as csv files for statistical analysis.

### Blood sampling and CORT quantification

Thirty and sixty minutes after the end of the post-training infusion, 500 μL of blood was collected from each rat through the tail-clip method. The most distal portion of the tail (approximately 1 mm) was removed with a scalpel blade; the tail was then gently manipulated so that blood accumulated at its tip and was collected into EDTA-coated tubes. The second collection, at 60 min, did not require a new incision: the cicatricial exudate was removed and the tail was manipulated as described above. Topical lidocaine and rifamycin spray were applied after the second collection to minimize discomfort. Blood samples were centrifuged at 2300 rpm for 20 min at 4 °C, and the resulting plasma was stored at −20 °C until hormone quantification.

Plasma CORT was measured by ELISA using the Corticosterone Enzyme Immunoassay Kit (Arbor Assays LLC, MI, USA), according to the manufacturer’s instructions. We excluded a subset of blood samples from CORT quantification (VEH n=3, MIF n=3) due to hemolysis or insufficient sample volume for accurate ELISA measurement. Optical densities were read at 450 nm using an Epoch spectrophotometer (BioTek Instruments, Inc.), and sample concentrations were calculated from the standard curve generated for each assay. All samples were analyzed in duplicate; the mean intra-assay coefficient of variation was 6.2 ± 0.9% and the inter-assay coefficient was 6.4%.

### Perfusion and immunohistochemistry

Ninety minutes following the end of the test session, rats were deeply anesthetized with 30% urethane and then perfused transcardially with 100 mL saline followed by 500 mL of ice-cold 4% paraformaldehyde (PFA) dissolved in phosphate buffer saline. The brains were removed, fixed for at least 2 h in PFA, then transferred to 30% sucrose solution and stored at 4 °C. After dehydration, brains were frozen in isopentane on dry ice and stored at −80 °C. Frozen coronal sections (40 μm) were cut in a Leica SM2010 R sliding microtome with dry ice and stored in 5 serial sets. One set was collected on glass slides and Nissl-stained for cytoarchitectural delineation of brain areas; the remaining sets were used for c-Fos immunolabeling.

Each immunohistochemistry assay included sections from all experimental groups and both timepoints, including homecage animals, to control for inter-assay variability. Free-floating sections were incubated at 4 °C for 72 h with a rabbit anti-c-Fos polyclonal antibody (1:15,000, abcam, ab190289) in 2% normal goat serum, 0.3% Triton X-100, and 0.02 M KPBS. After washes, sections were incubated with a biotinylated goat anti-rabbit secondary antibody (1:200, Vector Laboratories, Cat# BA-1000, RRID: AB_2313606) in 0.02 M KPBS and 0.3% Triton X-100 for 90 min at room temperature, followed by incubation in avidin-biotin-HRP complex solution (VectaStain Elite ABC Kit, Vector Laboratories, Cat# PK-6100). Peroxidase activity was visualized using the chromogen diaminobenzidine 3,3-tetrachloride (DAB Substrate Kit, Vector Laboratories, Cat# SK-4100). Sections were mounted on gel-coated slides, left to dry for at least 48 h, diaphanized, and coverslipped with DPX mountant medium (Sigma 06522) and left to dry for at least one week.

### Cell counting

c-Fos expression was analyzed in 12 regions of interest (ROIs): dorsal and ventral hippocampal subregions (dDG, dCA3, dCA1, vDG, vCA3, vCA1), amygdalar nuclei (BLA, CeA), and cortical regions (ACC, PrL, aIC, aRSC). Images from each ROI were acquired at 20x magnification using a Leica DM5500 microscope in brightfield mode (Table 1), in 24-bit RGB format at 2048 x 1536 pixel resolution (0.51 μm/pixel). The anatomical delimitation of each ROI was based on the Paxinos and Watson rat brain atlas (Paxinos & Watson, 2007) adjacent Nissl-stained sections.

**Table 1.** Description of microscopy for each region of interest (ROI)

| Region | Bregma Range | Number of Bilateral Frames per ROI | Median Bilateral Sections per Animal | Mean c-Fos Density Homecage Group |
|---|---|---|---|---|
| **dDG** | [-2.64, -3.60] | 2 | 3 | 69.71 |
| **dCA3** | [-2.64, -3.60] | 1 | 3 | 47.97 |
| **dCA1** | [-2.64, -3.60] | 2 | 3 | 88.01 |
| **PrL** | [3.24, 2.28] | 1 | 2 | 393.80 |
| **BLA** | [-1.92, -3.36] | 1 | 3 | 157.72 |
| **CeA** | [-1.92, -3.12] | 1 | 2.5 | 226.83 |
| **aRSC** | [-2.64, -3.60] | 1 | 3 | 1037.70 |
| **ACC** | [3.24, 2.28] | 1 | 2.5 | 409.15 |
| **aIC** | [3.24, 1.44] | 1 | 3 | 263.41 |
| **vDG** | [-4.80, -5.88] | 2 | 1 | 65.50 |
| **vCA3** | [-4.80, -5.88] | 1 | 1 | 44.00 |
| **vCA1** | [-4.80, -5.88] | 2 | 1 | 133.74 |

*Note: All images were acquired using the same microscope (Leica DM5500), with a 20× objective, an image size of 2048 x 1536 pixels, and a resolution of 0.51 µm per pixel. Frames were acquired from a range of bregma coordinates extending from the most anterior to the most posterior position for each brain region (column "Bregma Range"). Brain regions larger than the microscope's standard field of view were captured using more than one frame (column "Number of bilateral frames per ROI"). The column "Median bilateral sections per animal" shows the median number of bilateral sections per animal, considering all experimental groups.*

Cells were counted automatically using ImageJ (v1.52a, NIH). Images were converted to 8-bit grayscale and a contrast threshold was applied to binarize the signal; immunoreactive cells were then identified using the particle analysis tool with filters for size, circularity, and contrast. The intensity threshold was adjusted according to ROI- and assay-specific background staining, within a predefined range (lower threshold: 115–145; upper threshold: 255). For each ROI within an assay, the threshold was selected based on representative images from multiple animals and then applied consistently across the corresponding images. All experimenters were blinded to group assignment. Cell density was calculated by dividing the number of immunoreactive cells by the total image area. The mean c-Fos density per animal was normalized to the average density of homecage rats euthanized on the same days, yielding normalized c-Fos expression as the experimental unit. Counts from sections with tissue damage were excluded from the analysis.

### Behavioral correlation analysis and overview of computational workflow

First, we calculated the relationships among c-Fos expression levels of 12 brain regions (dDG, dCA3, dCA1, vDG, vCA3, vCA1, BLA, CeA, PrL, ACC, aIC, aRSC), behavioral variables and CORT measurements separately for VEH and MIF groups and timepoints (2 or 14 days) using Spearman correlations (’behavioral_correlations.ipynb’). Then, for c-Fos co-activation analyses, we considered each animal as one observation and each brain region as one variable. Within each experimental group, pairwise Spearman correlation coefficients were computed across animals for all pairs of brain regions, generating correlation matrices (’network_complete.py’). An optimal threshold for graph analysis was defined after label permutation (’permutation_analysis.py’). These correlation matrices were used to construct thresholded functional co-activation graphs, calculate graph-theoretical metrics, identify community structure and hub classifications, and compare correlation matrices between groups. Lastly, we compared c-Fos co-activation differences within and between predefined anatomical subnetworks among the experimental groups (’subnetworks.py’).

### Threshold determination by label permutation

To determine which coefficient threshold to use for functional coactivity graph analysis, we estimated a null distribution of Spearman correlations for graph binarization using the real dataset of c-Fos cell counting, following standard permutation-testing principles for empirical threshold estimation (Yu & Hutson, 2024). First, we computed real correlation matrices separately for the four groups formed by timepoint and treatment: 2 days VEH, 2 days MIF, 14 days VEH, and 14 days MIF. For each group, we calculated Spearman correlations with pairwise deletion of missing values. To avoid duplicate pairs, we extracted each unique correlation from the matrices and pooled the values as the real correlation. Then, to create the null distribution, we permuted across animals the treatment and timepoint labels while preserving the observed joint label structure and group sample sizes. For each permutation (n = 10,000, random seed 42), we recalculated group-wise correlation matrices for the four shuffled groups keeping the original sample size and pooling each unique pairwise-correlation. We summarized the null distribution by the 25th (rho = -0.0545), 50th (rho = 0.2667), 70th (rho = 0.4909), 75th (rho = 0.5476), 90th (rho = 0.7333), 95th (rho = 0.8155), and 99th (rho = 0.9167) percentiles. We selected the 70th percentile of the null distribution as the graph binarization threshold. This choice reflects a balance between two competing constraints inherent to small functional co-activation networks: retain enough edges for graph-theoretic analysis and the requirement that surviving edges exceed chance-level correlations. At more conservative thresholds (90th and 95th percentiles), the resulting graphs contained too few edges to yield connected networks across all 4 experimental groups, making it impossible to compute metrics dependent on path-length such as global efficiency (Supplementary Figure 2C). The 70th percentile represents the value above which the real correlation distribution is reliably higher relative to the null distribution, as confirmed by bootstrap resampling. In each bootstrap iteration (n = 10,000, random seed 42), null correlations were sampled with replacement to match the number of real correlations (N = 264) and the 2 groups means were compared using the Kolmogorov-Smirnov test. The distribution of p-values for K-S tests was then plotted and 89.4% of them were below the significance threshold of 0.05 (with mean p-value = 0.0134, Supplementary Figure 2E), confirming that the real distribution had significantly higher average value compared to the null distribution.

### c-Fos co-activity matrix construction

For each group, we calculated pairwise Spearman correlations for every pair of brain regions (Coelho et al., 2018). We then compared these group correlation matrices using Fisher r-to-z transformation. Correlation coefficients were clipped to the interval -0.9999 to 0.9999 to avoid infinite transformed values, and then transformed as:

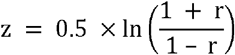

where z is the Fisher z value and r is the rho coefficient.

For each pair of regions, the difference between transformed correlations from the two groups (Z^1^ and Z^2^) was divided by the standard error (SE):

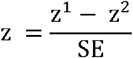

and SE was defined as:

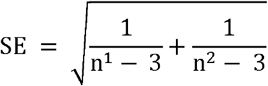

where n^1^ and n^2^ are the group sample sizes.

Two-tailed p values were calculated from the standard normal distribution as:

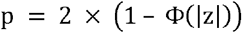

where is the cumulative distribution function of the standard normal and is the absolute value of the observed test statistic.

Significant pairwise group differences were defined at p < 0.05. To robustly control for multiple comparisons, we followed the previous step with an empirical permutation validation. For each brain-region pair, the observed absolute difference in Spearman correlation between groups was calculated (i.e., 14 days MIF dCA1-dCA3 minus 2 days MIF dCA1-dCA3). Animal-level rows from both groups were pooled, randomly permuted (n = 1,000, random state 42), and split into pseudo-groups matching the original group sizes. For each permutation, Spearman correlations were recalculated for the two pseudo-groups and the absolute difference between pseudo-group correlations was stored. Empirical p values were calculated as:

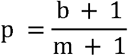

where b is the number of permuted absolute differences greater than or equal to the observed absolute difference and m is the number of valid permutations.

### Thresholded graph construction

To investigate differences between functional c-Fos coactivity between different groups, we constructed undirected weighted graphs from the previously described correlation matrices using NetworkX, as described elsewhere (dos Santos Corrêa et al., 2026). We defined a graph structure G as the tuple of vertices and edges (V, E), where V represents brain regions, E represents functionally connected pairs from the adjacency matrix A, and W is the weighted adjacency matrix. This weighted, undirected graph can be expressed mathematically as:

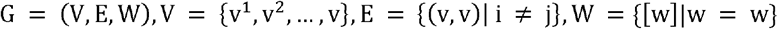

Nodes represented brain regions and edges represented above threshold positive coefficients. Edge weights were stored as the absolute correlation coefficient for weighted graph metrics. Community structure was detected for each graph using Louvain community detection with absolute edge weights, resolution of 0.6, and random state set to 42.

For each group-level graph, we calculated five global network metrics: mean degree, mean strength, weighted average clustering coefficient, global efficiency, and modularity. Mean degree was calculated as the average number of edges per node. Mean strength was calculated as the average sum of absolute edge weights connected to each node across all nodes. Weighted average clustering was calculated using the function ‘networkx.average_clustering()’. Global efficiency was calculated using the function ‘networkx.global_efficiency()’. Modularity was obtained from the community detection step using the function ‘community_louvain.modularity()’.

Next, we classified nodes using within-module degree z-scores and participation coefficients calculated from the community structure, following a previously reported method (Meunier et al., 2010; Santos et al., 2023). For each node, the within-module z-score (z) was quantified as:

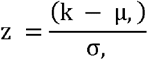

Where k is the within-module degree of node I (number of edges connecting node i to other nodes within its own community), µ, is the mean within-module degree of all nodes in community s and σ, is the standard deviation of within-module degree in the community s.

The participation coefficient (PC) was calculated as:

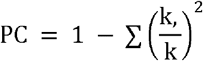

Where k,. is the number of edges from node i to nodes in the community m, k is the total degree of node I, and the sum runs over all communities m in the network.

We next assigned hub categories. Due to the small number of nodes (12), we adapted the thresholding method to our data. Nodes with within-module z-score >= 0.16 were classified as provincial hubs when participation coefficient was less than 0.30 and connector hubs when participation coefficient was at least 0.30 and less than 0.75. Nodes with within-module z-score below 0.16 were classified as non-hub connectors when participation coefficient was at least 0.30. If nodes did not fall under these categories, they were classified as ultra-peripheral. Additionally, to describe how connections were spread across brain regions, we calculated the degree of each node in each treatment and timepoint group. Within the graph for each group, we ordered the 12 regions from highest to lowest degree. We colored each region by its Louvain community assignment, so that degree ranking could be read alongside community structure.

### Subnetwork-level co-activation analysis

To further investigate the differences observed in graph analysis, we defined subnetworks according to anatomical classes of brain regions. Dorsal hippocampus (dHPC) contained dDG, dCA3, and dCA1; ventral hippocampus (vHPC) contained vDG, vCA3, and vCA1; total hippocampus (HPC) combined dHPC and vHPC; amygdala contained BLA and CeA; and cortex contained ACC, PrL, aIC, and aRSC. We assigned each pairwise functional connectivity edge to one of two resting-state networks: a default mode network (DMN)-like network or a salience network (SN)-like network, based on the established network membership of its constituent regions of interest (ROIs). We assigned edges involving the anterior cingulate cortex (ACC), prelimbic cortex (PrL), and anterior retrosplenial cortex (aRSC) to the DMN, consistent with the role of these regions as core hub nodes of the rodent DMN. We assigned edges involving the anterior insular cortex (aIC), basolateral amygdala (BLA), and central amygdala (CeA) to the SN, consistent with the canonical insular-amygdala axis of the SN. We classified each edge according to the network affiliation of its region pair rather than a single fixed region list, so a given region contributed edges to both networks depending on its correlation partner. For example, we labeled BLA-ACC as a DMN edge but labeled BLA-aIC as an SN edge, because we defined network membership at the level of the edge, not the node. We excluded the aIC-aRSC edge from both subnetworks because its constituent ROIs belong one each to the DMN-like seed set (aRSC) and the SN-like seed set (aIC), making its network assignment ambiguous under our edge-level classification scheme. We also excluded two edges from the subnetwork-level analysis: BLA-aRSC and dCA1-aIC. We excluded these edges because they connected a node from one network’s canonical seed set to a node from the other network’s non-seed but functionally affiliated regions.

After generating pair lists for the subnetworks (hippocampus-within: within HPC, hippocampus-amygdala: HPC-Amy, hippocampus-cortex: HPC-CTX, and amygdala-cortex: Amy-CTX) or resting state networks, we transformed each pairwise correlation using z = arctanh(r), with correlations clipped to - 0.9999 to 0.9999 before transformation. For each group and subnetwork, we calculated the mean Fisher z value across edges, then we performed statistical tests using these values. Z values were then back-transformed to rho using the hyperbolic tangent for interpretation and plotting.

## QUANTIFICATION AND STATISTICAL ANALYSIS

### General statistical approach

Unless otherwise stated, analyses were performed in Python (v. 3.11.8) or JAMOVI (v.2.7.39) (The jamovi Project, 2020). Figure panels were plotted using Graphpad Prism (v 11.0) or Python. Data was tested for normality, homogeneity and sphericity before all parametric tests that required them. Spearman correlations were used for correlation tests to reduce reliance on linearity and normality assumptions. Missing values were handled by pairwise deletion for correlation analyses, when applicable. For c-Fos network analyses, region-pair correlations required at least six valid paired animal observations.

We quantified freezing, rearing, grooming, and sniffing separately in the original conditioned context (A) and the novel context (B). We expressed freezing as a percentage of session time and rearing, grooming, and sniffing as total duration per session. For analyses of overall behavioral magnitude, we calculated the mean across contexts as (XA+XB)/2, where XA and XB denote the corresponding behavioral measurements in contexts A and B. We calculated the freezing discrimination index as (FA−FB)/(FA+FB). Where FA refers to freezing in the original context and FB freezing in the novel context. Positive values indicated greater freezing in the conditioned context, zero indicated equal freezing across contexts, and negative values indicated greater freezing in the novel context. For rearing, we calculated novel-context preference as (RB −R A )/(R A +R B). RB refers to total rearing in the novel context and RA, rearing in the original context.

We summarized post-training corticosterone (CORT) concentrations using a decay ratio and the area under the curve (AUC). We calculated the decay ratio as (CORT30−CORT60)/(CORT30+CORT60), using blood samples collected 30 and 60 minutes after the end of contextual fear conditioning. Higher values indicated a larger relative decline between measurements, zero indicated equal concentrations, and negative values indicated an increase. We calculated CORT AUC using the trapezoidal rule: [(CORT 30 +CORT 60 )/2]×30. We expressed AUC in (ng/mL)min. to quantify integrated corticosterone concentrations over the sampled interval.

### Statistical analysis of Behavioral and Endocrine measures

Statistical tests are indicated in the figure legends. To evaluate statistical significance from most behavioral analysis, data from **Figures 1 C-D; Fig. 2 B-C/F-G** were subjected to Mixed Model ANOVAs followed by Tukey post hoc tests. Data from **Figure 1 E; Supplementary Figure 1 A** was subjected to Welch’s t tests. To analyze the composition of behavioral bouts (Figure S1B), we built contingency tables of behavioral category (freezing, rearing, grooming, sniffing) by context (original, novel), stratified by treatment, separately for each retention interval. We tested the context-behavior association across treatments with a generalized Cochran-Mantel-Haenszel test. To test whether this association differed between treatments, we compared log-linear models with and without the Treatment x Context x Behavior term using a likelihood-ratio G² test. Within each treatment, we tested the context-behavior association with Pearson’s χ² tests. We then identified the behavioral categories driving significant associations from adjusted standardized residuals (|z| > 1.96; two-sided p < 0.05). These residuals indicate cellwise departures from independence and do not compare treatments. We ran these analyses with the Chi-Squared Analysis Tools module (v1.2.1) in JAMOVI. Spearman correlations with Benjamini-Hochberg false discovery rate were calculated for data in **Figures 1 F-G; Fig. 2 E/I; and Fig. 5 B/D-G**. We present data as median, interquartile range, and minimum/maximum values, except for time-course data (**Figure 2B, F, J; Figure S1C, F**), which we present as mean ± SEM and the threshold for significance was at p < 0.05, unless noted otherwise.

We employed a general linear mixed model (LMM) approach for the minute-by-minute freezing and rearing duration analysis. The LMM was chosen for its ability to handle both fixed and random effects, making it particularly suitable for our repeated measures design and hierarchical data structure (de Melo et al., 2022). The model was implemented using the General Analyses for Linear Models module (GAMLj3, v3.7.2) in JAMOVI. We applied two factor coding schemes to address specific research questions and facilitate interpretable comparisons between repeated measures and conditions, with individual rats and the interaction between context * rats as the random factors. Simple factor coding was always employed for comparisons between the Treatment and Context factors, and their interactions. For the Minutes factor and all its interactions, we used Helmert coding which compared each level to the mean of all next levels, allowing us to investigate how each condition differed from the average of the conditions that followed it (e.g., minute 1 vs minute 2-4) (**Figure 2J, Supplementary Figure 1 C/F**). Multiple comparisons were corrected using the Tukey method where needed. We analyzed grooming duration (**Figure S1D, G**) with a generalized linear mixed model (GLMM) with a Gamma distribution and an identity link, Treatment and Context as fixed effects, and animal as a random intercept. We chose this model because grooming durations were positively skewed. Because the Gamma distribution requires strictly positive values, we added 1s to every data point before model fitting. We tested fixed effects with omnibus χ² tests using GAMLj3 in JAMOVI.

We used principal component analysis (PCA) to summarize how defensive behavior and post-training corticosterone (CORT) covaried. We entered six variables per animal. The first five came from the retrieval tests in the training context (A) and the novel context (B): mean freezing, freezing discrimination index, mean rearing, rearing discrimination index, grooming context difference. The sixth was CORT Decay Ratio. Six animals from the 14-day cohort had no CORT Decay Ratio values, so for each of them we filled in the value of a randomly chosen animal from the same treatment group and retention interval. We standardized each variable across all animals and ran the PCA on the standardized data **(Figure 2K)**. To decide how many components to keep, we used parallel analysis. We shuffled each variable independently across animals 5000 times and kept a component only if its eigenvalue was above the 95^th^ percentile of the shuffled eigenvalues for that component. Only the first principal component (PC1) passed this test **(Figure S1I)**. We oriented PC1 so that mean freezing had a positive coefficient. We then compared PC1 scores with a two-way ANOVA with Treatment and Retention interval as factors (**Figure 2L**).

### Statistical analysis of c-Fos expression levels

For each brain region, c-Fos expression levels were compared between 2 and 14 days for either MIF or VEH groups using Welch’s two-sample t test. Between-group effect size was calculated as Cohen’s d value. One-sample t tests compared each group against the average of c-Fos expression from a homecage group of animals, which was equal to 1.0. Benjamini-Hochberg false discovery rate correction was applied separately to the family of between-group tests, the family of VEH-versus-homecage tests, and the family of MIF-versus-homecage tests (**Figure 5 A/C, Supplementary Figure 6 A-B**). For the correlation matrices, we used Spearman correlations as explained before (**Supplementary Figure 3 A-B and Supplementary Figure 4 A-B**) For the Fisher matrix comparisons and permutation validation of individual correlation differences significance threshold was set at p < 0.05 (**Figure 4 F/H; Supplementary Figure 3 C and Supplementary Figure 4C**). For the subnetwork-level or resting state networks co-activation analysis, we used bootstrapped 1-way ANOVAs with the Treatment|Timepoint as factor with additional Bonferroni-type post hoc tests (**Figure 3 C; Fig. 4 B-E/G/I**). For bootstrapping, we used a modified one-step estimator and 2000 samples with Mahalanobis distances. For all analyses data are presented as median ± max/min and the threshold for significance was at p < 0.05, unless noted.

### Animal-level permutation testing of network metrics

We evaluated group differences in global network metrics using animal-level permutation tests following a previously reported method (Simpson et al., 2013). For each metric, observed group-level metric values were calculated from the original VEH and MIF-treated rats, and the observed difference was calculated as MIF minus VEH (i.e.: MIF mean degree minus VEH mean degree). For each permutation (n = 10,000; random seed = 42), animals from both groups were pooled and randomly reassigned into pseudo-groups preserving original group sizes. The full graph reconstruction and metric calculation were repeated using the same 70th percentile coefficient extracted from the null distribution described on subitem “Threshold determination by label permutation”, meaning that the significance of reported group differences does not depend on the absolute threshold value but on whether observed network differences exceed those expected by chance under the same binarization rule. Finally, two-sided empirical p-values were calculated as the proportion of absolute permuted differences greater than or equal to the absolute observed difference. We considered the observed difference statistically significant when the empirical p-value was < 0.05, indicating that the observed difference was unlikely to have arisen under the null distribution generated by permutation (**Figure 3 J-K/M-N; Supplementary Figure 3 D-F/J-L**).

### Python packages

The code was implemented in Python and makes use of the following packages: ‘numpy’ (v1.26.4), ‘scipy’ (v1.11.4) and ‘pandas’ (v2.1.4); ‘Matplotlib’ (v3.9.2), ‘seaborn’ (v0.13.2) for data visualization; ‘networkx’ (ver. 3.1), ‘igraph’ (ver. 0.11.8) for Network analysis.

