## Supplemental Figures for "Prefrontal cortex glucocorticoid receptors during fear memory consolidation shift the balance between salience and default-mode networks at retrieval in rats"

**SUPPLEMENTARY INFORMATION**

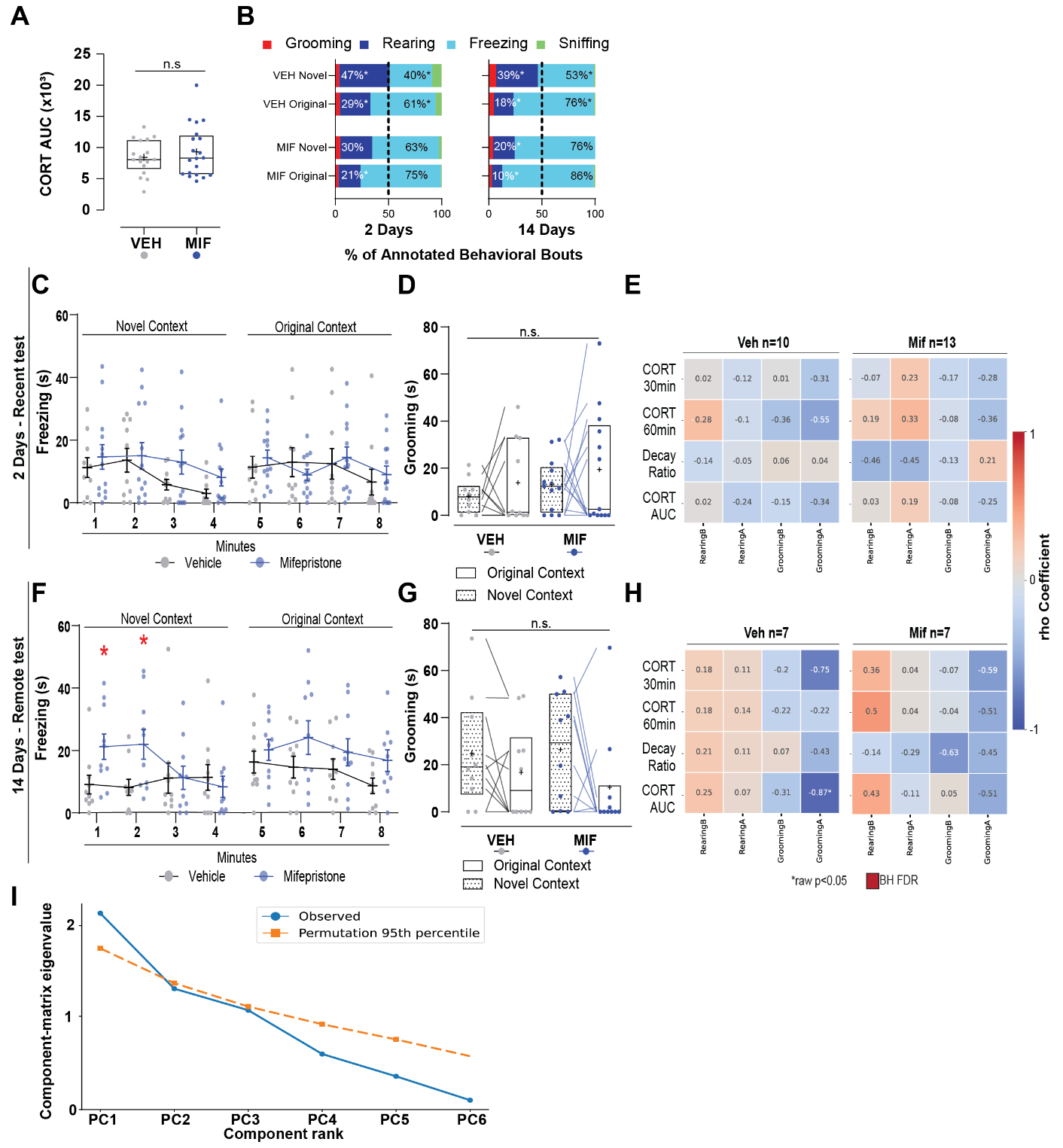

**Figure S1, related to Figures 1 and 2: Supplemental behavioral and corticosterone analyses and principal component retention**

**(A**) Median area under the curve (AUC) of CORT concentrations over the 30–60 min post-training interval for VEH- and MIF-treated rats as an integrated measure of stress hormone release. Boxplots showing median, interquartile intervals and maximum/minimum datapoints summarize data distribution. The plus sign reflects the mean for each subset of data. Two samples Student’s t test: t(35) = 0.75, p = 0.46, d = 0.25.

**(B)** Distribution of annotated behavioral bouts at recent (2 days) and remote (14 days) memory tests. Stacked bars show the percentage of grooming, rearing, freezing, and sniffing bouts in the original and novel contexts for VEH and MIF-treated rats. At 2 days, context and behavioral category showed a significant association (generalized Cochran-Mantel-Haenszel test, 𝜒2(3)=44.1, 𝑝<0.001), with no evidence that this association differed between treatments (log-linear likelihood-ratio test, 𝐺2(3)=3.66, 𝑝=0.30). At 14 days, context-behavior associations differed between treatments (𝐺2(3)=7.98, 𝑝=0.046). Both groups showed significant associations (VEH: 𝜒2(3)=38.9; MIF: 𝜒2(3)=20.5; both 𝑝<0.001). Asterisks mark significant cellwise departures from independence for rearing and freezing (adjusted standardized residuals, ∣𝑧∣>1.96; two-sided 𝑝<0.05), not between-treatment comparisons.

**(C)** Mean freezing times (s) ± S.E.M. across consecutive minutes in the novel (left) and original (right) contexts for VEH- and MIF-treated rats during the 2 days retrieval test. Linear Mixed Model: Context *F*(1,21) = 0.12, *p* = 0.73; Minutes *F*(3,126) = 8.03, *p* < 0.001; Treatment *F*(1,21) = 0.51, *p* = 0.48; interactions not significant (Random effects: Animal, Context|Animal).

**(D)** Mean grooming times (s) ± I.Q.R. Lines connect measurements from the same animal. We analyzed grooming duration using a generalized linear mixed-effects model with a Gamma distribution, an identity link, Treatment and Context as fixed effects, and animal as a random intercept. Neither Treatment (𝜒2(1)=1.94, 𝑝=0.164), Context (𝜒2(1)=1.55, 𝑝=0.213), nor their interaction (𝜒2(1)=0.054, 𝑝=0.816) significantly affected grooming duration (𝑛=23). n.s., not significant.

**(E)** Heatmaps of Spearman correlations between corticosterone (CORT) measures and behavioral measures at the recent memory test (2 days), separately for VEH and MIF-treated rats. CORT measures include concentrations at 30 and 60 min, the decay ratio, and the area under the curve (AUC). Cell values indicate Spearman’s 𝜌; colors represent correlation direction and magnitude. Asterisks indicate unadjusted 𝑝<0.05; thick outlines indicate significance after Benjamini–Hochberg false discovery rate correction (𝑞<0.05). VEH, 𝑛=10; MIF, 𝑛=13.

**(F)** Mean freezing times (s) ± S.E.M. across consecutive minutes in the novel (left) and original (right) contexts for VEH- and MIF-treated rats during the 14 days retrieval. Linear Mixed Model: Context *F*(1,18) = 2.18, *p* = 0.157; Minutes *F*(3,108) = 4.52, *p* = 0.005; Treatment *F*(1,18) = 3.18, *p* = 0.09; Minute*Treatment *F*(3,108) = 2.94, *p* = 0.037; Minute*Context*Treatment *F*(3,108) = 2.96, *p* = 0.035; other interactions not significant (Random effects: Animal, Context|Animal). *p < 0.05 when compared to the following minutes only in the MIF-treated group.

**(G)** Total grooming duration in the novel and original contexts at 14 days, plotted as in (G). Generalized linear mixed model: Treatment (𝑝=0.154), Context (𝑝=0.167), or their interaction (𝑝=0.572; 𝑛=20). n.s., not significant.

**(H)** Heatmaps of Spearman correlations between CORT measures and behavioral measures at the remote memory test (14 days), separately for VEH- and MIF-treated rats. Colors, cell values, and significance annotations follow (E). 𝑛=7 per treatment.

**(I)** Parallel analysis of principal-component eigenvalues. Blue circles show observed correlation-matrix eigenvalues; orange squares show the rank-specific 95th percentile of null eigenvalues from 5,000 permutations, each independently shuffling animal values within each variable. Only the first principal component (PC1) exceeded its permutation threshold, supporting retention of one component.

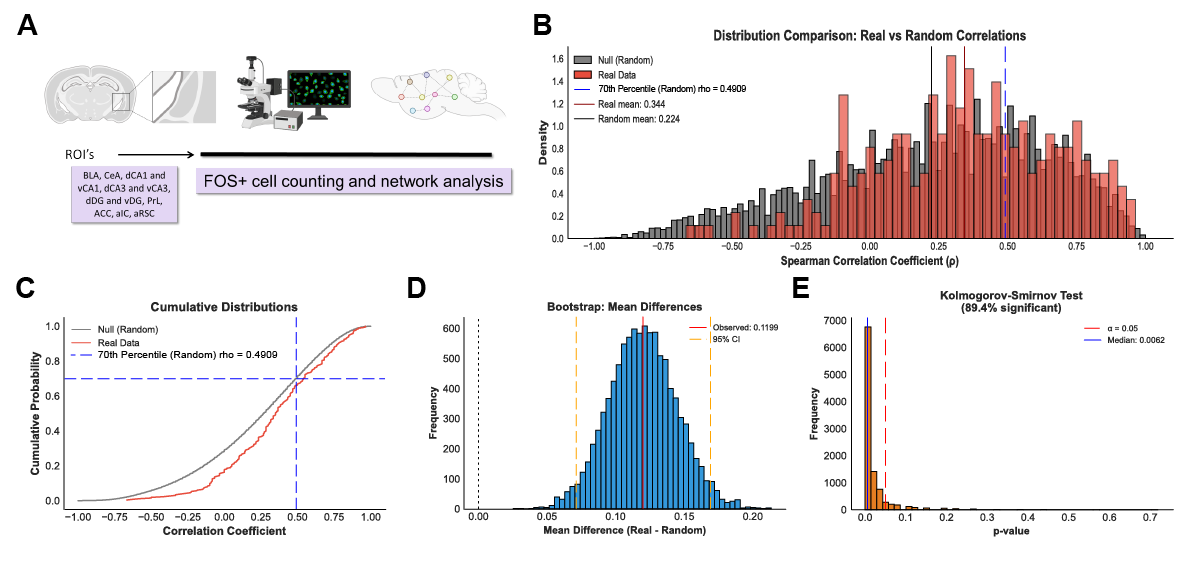

**Figure S2, related to Figures 3 and 4: Empirical threshold determination for functional co-activation graphs via label permutation**

**(A)** Workflow schematic: brain regions of interest (BLA, CeA, dCA1, vCA1, dCA3, vCA3, dDG, vDG, PrL, ACC, aIC, aRSC) underwent c-Fos immunolabeling, brightfield microscopy imaging, and Fos+ cell counting to build region-by-region functional co-activation networks.

**(B)** Density histograms comparing the real pairwise Spearman correlation coefficients (red) against the null distribution generated by label permutation (gray; n = 10,000 permutations, random seed 42, treatment and timepoint labels shuffled across animals while preserving group sizes). The dashed blue line marks the 70th percentile of the null distribution (rho = 0.4909), which was chosen a priori as the threshold for graph binarization; solid lines indicate the real (mean = 0.344) and random (mean = 0.224) distribution means.

**(C)** Cumulative distribution functions of the real and null correlation coefficients, with the horizontal and vertical dashed blue lines marking the 70th-percentile threshold (rho = 0.4909) used for thresholded graph construction.

**(D)** Bootstrap distribution (n = 10,000 iterations, random seed 42) of the mean difference between real and null correlation coefficients (real correlations sampled with replacement to match N = 264); the red line marks the observed mean difference (0.1199) and orange dashed lines mark the 95% confidence interval, which excludes zero (dotted black line).

**(E)** Distribution of 𝑝-values from Kolmogorov-Smirnov tests comparing resampled observed and label-permuted correlation distributions. The red dashed line marks 𝑝=0.05, and the blue line marks the median 𝑝-value (0.0062). Of the resampling iterations, 89.4% yielded 𝑝<0.05.

.

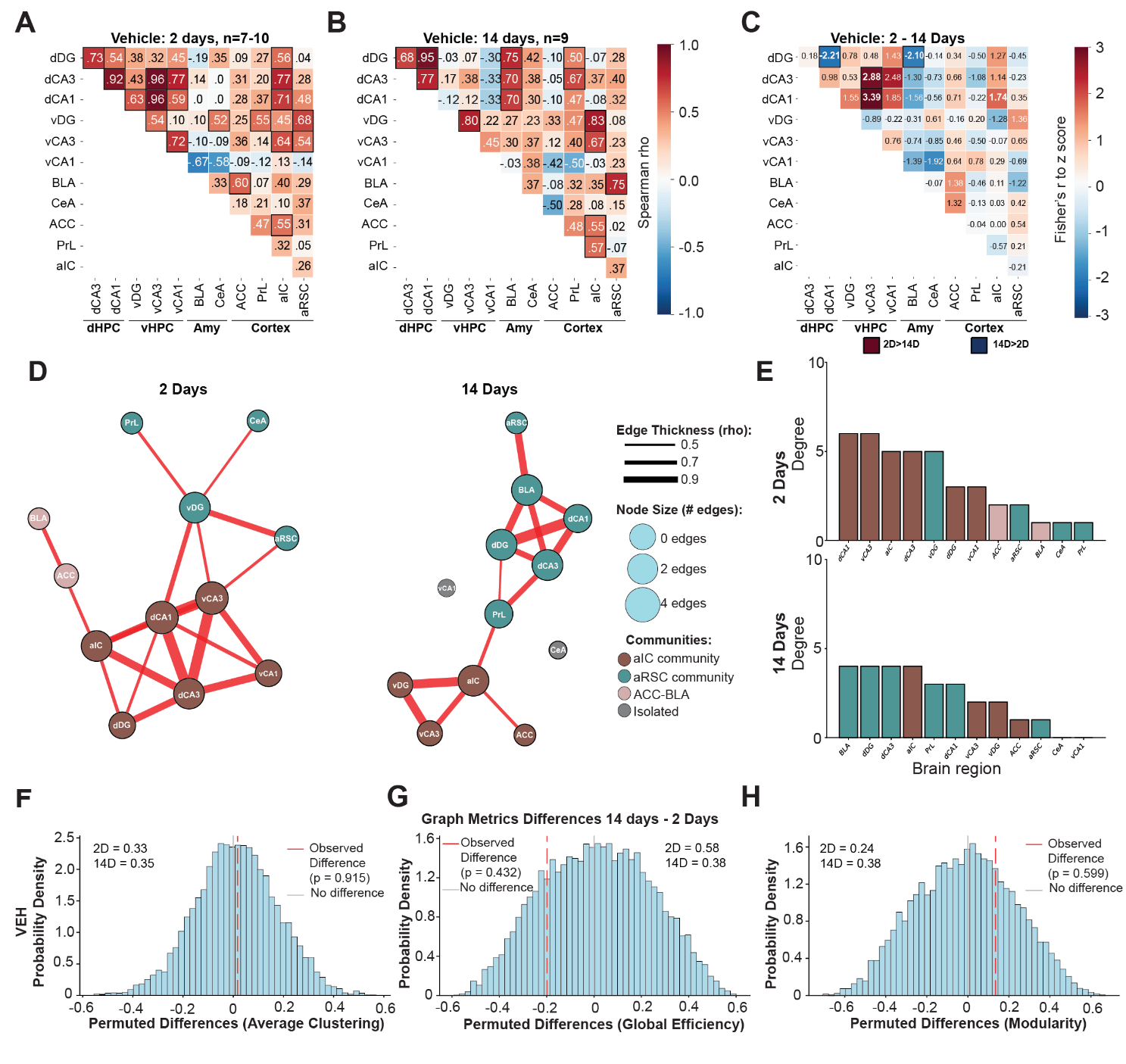

**Figure S3, related to Figure 3. C-Fos coactivation networks and graph analysis for VEH-treated animals at both timepoints**

**(A)** Regional c-Fos co-activation matrices in vehicle-treated rats at 2 days retrieval test. Heatmaps show pairwise Spearman correlation coefficients (rho) between c-Fos expression in dorsal hippocampus (dDG, dCA3, dCA1), ventral hippocampus (vDG, vCA3, vCA1), amygdala (CeA, BLA), and cortical regions (ACC, PrL, aIC, RSC. n = 7-10).

**(B)** Spearman correlation matrix for VEH animals 14 days after conditioning (n = 9), plotted as in (A).

**(C)** Fisher z-score matrix showing the difference in c-Fos co-activation between 14 days and 2 days within the VEH group; boxed values with asterisks indicate significant pairwise differences (Fisher r-to-z transformation, two-tailed test, p < 0.05, validated by permutation testing).

**(D)** Co-activation networks for VEH animals at 2 days (left) and 14 days (right). Nodes represent brain regions, and edges represent positive, suprathreshold Spearman correlations between regional c-Fos counts across animals. Edge thickness scales with the correlation coefficient (rho), and node sizes scales with degree (number of connections). Node colors indicate community membership; gray identifies isolated nodes.

**(E)** Regional node degrees for VEH networks at 2 days (top) and 14 days (bottom), ordered from highest to lowest. Bar colors match community assignment in (D).

**(F-H)** Animal-level permutation tests (n = 10,000 permutations, random seed 42) comparing global network metrics between 14-day and 2-day VEH graphs: average clustering coefficient (D), global efficiency (E), and modularity (F). Gray histograms show the null distribution of permuted differences; the red dashed line marks the observed 14D-2D difference, with empirical p-values reported in each panel and the corresponding group metric values annotated (2D and 14D).

Abbreviations: dDG, dorsal dentate gyrus; dCA3, dorsal CA3; dCA1, dorsal CA1; vDG, ventral dentate gyrus; vCA3, ventral CA3; vCA1, ventral CA1; BLA, basolateral amygdala; CeA, central amygdala; ACC, anterior cingulate cortex; PrL, prelimbic cortex; aIC, anterior insular cortex; aRSC, anterior retrosplenial cortex; dHPC, dorsal hippocampus; vHPC, ventral hippocampus; Amy, amygdala.

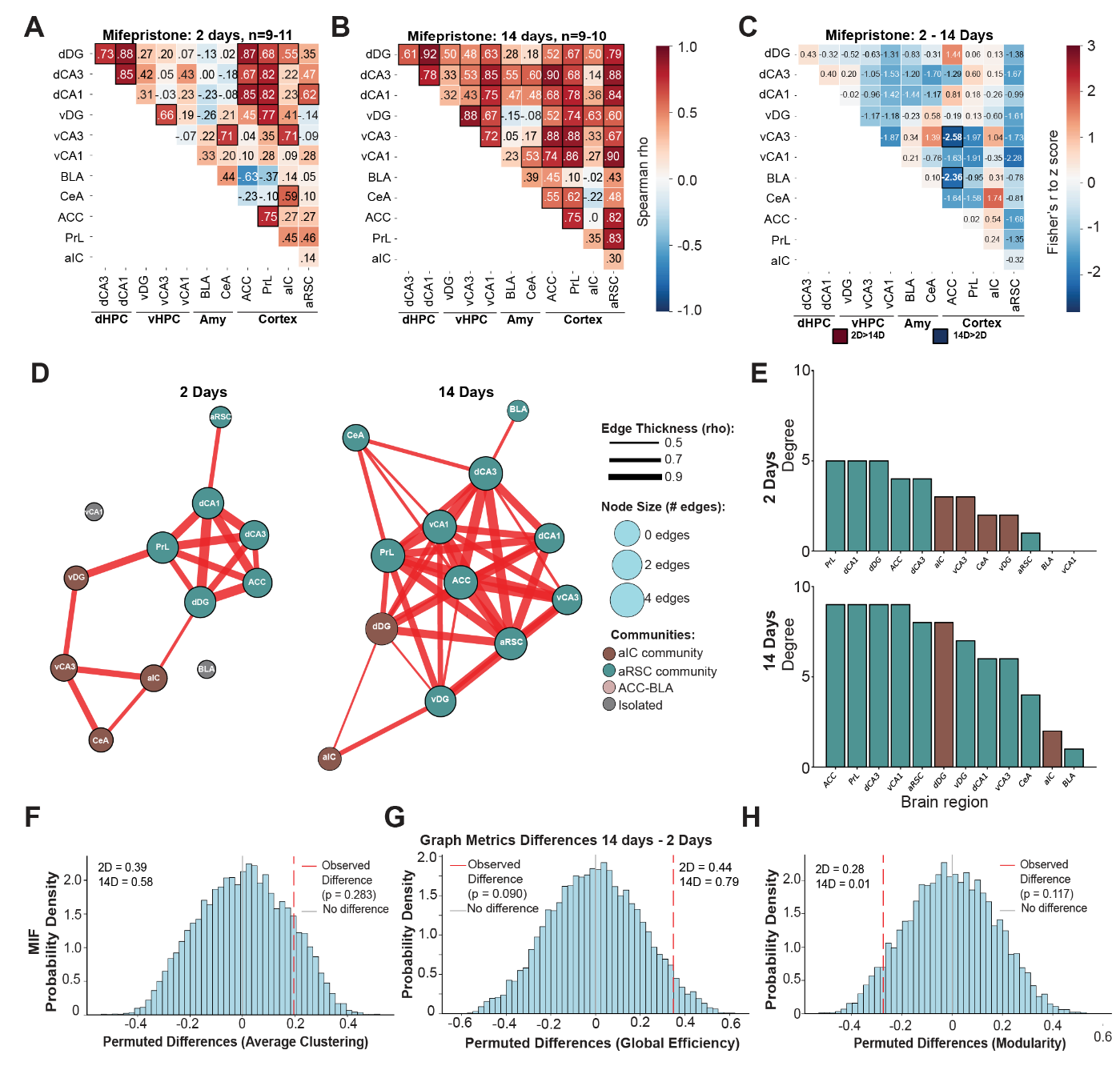

**Figure S4, related to Figure 3. C-Fos coactivation networks and graph analysis for MIF-treated animals at both timepoints**

**(A)** Spearman correlation matrix of c-Fos co-activation in mifepristone (MIF)-treated animals 2 days after conditioning (n = 9-11), plotted as in (Figure S3A).

**(B)** Spearman correlation matrix for MIF animals 14 days after conditioning (n = 9-10), plotted as in (Figure S3A).

**(C)** Fisher z-score matrix showing the difference in c-Fos co-activation between 14 days and 2 days within the MIF group, plotted as in (Figure S3C).

**(D)** Co-activation networks for MIF animals at 2 days (left) and 14 days (right), displayed as in (Figure S3D).

**(E)** Regional node degrees for MIF networks at 2 days (top) and 14 days (bottom), ordered from highest to lowest. Bar colors match community assignment in (D).

**(F-H)** Animal-level permutation tests comparing global network metrics between 14-day and 2-day MIF graphs: average clustering coefficient (F), global efficiency (G), and modularity (H), plotted as in (Figure S3F-H).

Abbreviations: dDG, dorsal dentate gyrus; dCA3, dorsal CA3; dCA1, dorsal CA1; vDG, ventral dentate gyrus; vCA3, ventral CA3; vCA1, ventral CA1; BLA, basolateral amygdala; CeA, central amygdala; ACC, anterior cingulate cortex; PrL, prelimbic cortex; aIC, anterior insular cortex; aRSC, anterior retrosplenial cortex; dHPC, dorsal hippocampus; vHPC, ventral hippocampus; Amy, amygdala.

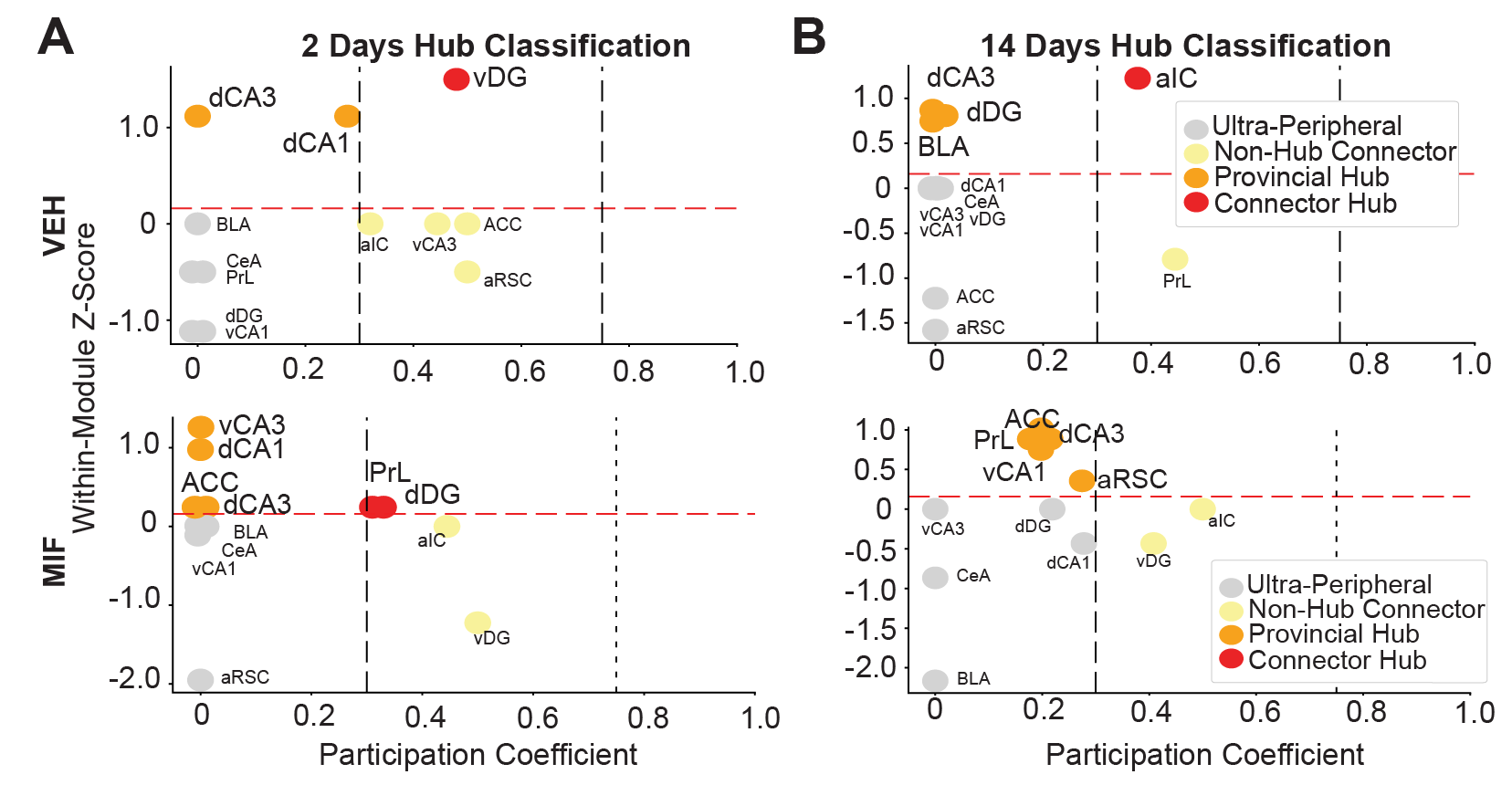

**Figure S5, related to Figure 4. Hub classification analysis**

**(A)** Hub classification for VEH (top) and MIF (bottom) networks at 2 days retrieval. Each point represents a brain region and plots its within-module degree z-score against its participation coefficient. Dashed lines indicate boundaries for ultra-peripheral (gray), non-hub connector (yellow), provincial hub (orange), and connector hub (red) categories.

**(B)** Hub classification for VEH (top) and MIF (bottom) networks at 14 days retrieval, displayed as in (A).

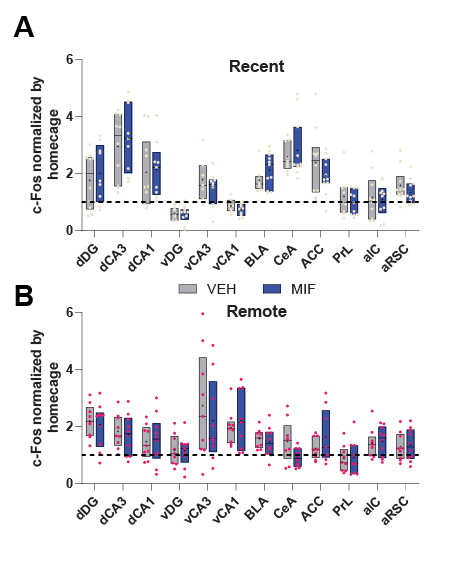

**Figure S6, related to Figure 5. Regional c-Fos expression relative to homecage baseline at recent and remote timepoints**

**(A)** c-Fos expression, normalized to a homecage control group (dashed line = 1.0), across 12 brain regions in VEH (gray) and MIF (blue) animals at the 2-day recent timepoint. Each dot (pink) represents one animal; box plots show median and interquartile range with "+" marking the mean. One-sample t tests compared each group against the homecage average, and Welch's two-sample t tests compared VEH vs. MIF within each region, with Benjamini-Hochberg false discovery rate correction applied separately within each test family.

**(B)** c-Fos expression normalized to homecage baseline across the same 12 brain regions in VEH (gray) and MIF (blue) animals at the 14-day remote timepoint, plotted and analyzed as in (A). Each dot (red) represents one animal.
